# Selection against dispersers: Sex, age and origin-dependent fitness differences in wild Atlantic salmon

**DOI:** 10.64898/2025.12.30.696970

**Authors:** Emilio Egal, Charles Perrier, Yann Czorlich, Geir H. Bolstad, Craig R. Primmer, Aurélie Manicki, Josu Elso, Guillaume Evanno, Mathieu Buoro

**Affiliations:** DECOD, INRAE, Institut Agro, IFREMER, Rennes, France; Management of Diadromous Fish in their Environment, OFB, INRAE, Université de Pau et des Pays de l’Adour, Institut Agro, Saint-Pée-sur Nivelle, France; CBGP, INRAe, CIRAD, IRD, Montpellier SupAgro, Université de Montpellier, Montpellier, France; Norwegian Institute for Nature Research (NINA), NO-7485 Trondheim, Norway; Organismal and Evolutionary Biology Research Program, Faculty of Biological and Environmental Sciences, University of Helsinki, Helsinki 00014, Finland; Université de Pau et des Pays de l’Adour, INRAE, ECOBIOP, Saint-Pée-sur-Nivelle, France; Área de Gestión Piscícola, Orekan Gestión Ambiental de Navarra, S.A., Pamplona/Iruña, Navarre, Spain

**Keywords:** Local adaptation, Dispersal costs, Reproductive success, Sex-biased dispersal, Pedigree analysis, Gene flow, Atlantic Salmon

## Abstract

Local adaptation is often portrayed as uniform fitness disadvantages of immigrants relative to residents. Yet dispersal costs vary with origin, sex, and life-history traits, shaping the balance between gene flow and adaptation. We quantified these heterogeneous costs by comparing the reproductive success of local and immigrant individuals using a 15-year genetic pedigree of Atlantic salmon (*Salmo salar*), including over 1,100 adults and 3,400 juveniles. Immigrants represented 19.5% of adults, showed male-biased dispersal, and averaged 15% lower reproductive success than locals. However, outcomes were strongly origin-dependent: immigrants from some populations suffered severe fitness costs while others matched or exceeded locals. Sex and life-history further modulated reproductive success: male immigrants underperformed local males, whereas multi-sea-winter females often equaled or outperformed female residents. Immigrant males partially compensated their fitness offset by acquiring more mates. These findings highlight that, while local adaptation promotes philopatry, immigrants*’* origin effects should be considered in gene-flow adaptation dynamics.

## Introduction

While immigration can introduce substantial demographic and genetic influences into local populations, selection against immigrants represents a key evolutionary force that can simultaneously restrict gene flow and promote local adaptation (Blanquart *et al*. 2013; Kawecki & Ebert 2004; Savolainen *et al*. 2007). Selection influences the reproductive success (RS) of immigrants and their descendants through multiple mechanisms (Nosil *et al*. 2005; Tobler *et al*. 2009), generating fitness costs that may extend beyond the initial dispersal event (Lenormand 2002). Ignoring the contribution of immigrants, and the mechanisms that facilitate or impede gene flow, can lead to misleading interpretations of the causes of change in local populations, particularly if immigrants differ in their life-history traits. Therefore, quantifying not only the number of immigrants but also their characteristics and relative reproductive success is essential for understanding the eco-evolutionary dynamics of populations.

Achieving this requires not only identifying dispersal events, i.e. any movement of individuals with potential consequences for gene flow across space (Ronce 2007), but also tracking their descendants. This is because the proportion of immigrants does not necessarily reflect the extent of gene flow, and vice versa (Lenormand 2002), as immigrants may experience pre- and/or postzygotic processes that either restrict or facilitate gene flow. Immigrants may face prezygotic barriers through natural selection (Lin *et al*. 2008; Matute *et al*. 2009; Nosil *et al*. 2005) or sexual selection, via behavioural incompatibilities, site familiarity, or mate choice (Higashi *et al*. 1999; Snowberg & Benkman 2009; Tobler *et al*. 2009). Immigrants and their descendants could also suffer from postzygotic effects including genetic incompatibilities and hybrid inviability (Nosil *et al*. 2005). Accordingly, F1 offspring of immigrant-local matings may make a limited genetic contribution to the local population if they inherit maladaptive alleles from immigrant parents, or a tendency to disperse that leads to their emigration soon after recruitment, thereby reducing effective gene flow (Doligez & Pärt 2008; Hansson *et al*. 2003). Critically, these costs may be cryptic in F1 generations but emerge in subsequent generations through increased expression of recessive deleterious alleles from both local and immigrant lineages and epistatic breakdown (Dickel *et al*. 2024; Hasselgren *et al*. 2024; Marr *et al*. 2002). Conversely, immigrants may also have higher fitness than resident individuals by exhibiting distinctive behaviours (boldness, aggressiveness) that enhance competitive ability for mates and resources (Clobert *et al*. 2009; Duckworth & Badyaev 2007) and also via reduction of inbreeding depression (Matthysen 2012). F1 offspring of immigrant-local matings could have a relatively higher fitness due to heterosis, increasing transgenerational gene flow (Hasselgren *et al*. 2018; Marr *et al*. 2002; Reid *et al*. 2024; Saatoglu *et al*. 2025).

Fitness of immigrants is increasingly recognized as being context-dependent, varying according to ecological and genetic differences (Clobert *et al*. 2009; Greenwood 1980; Hansson *et al*. 2004; Lin *et al*. 2008; Marr *et al*. 2002; Matute *et al*. 2009; Mobley *et al*. 2019; Reid *et al*. 2024; Richardson *et al*. 2014). These context-dependent effects are further complicated by fundamental differences in how selection operates between sexes. Female reproductive success (RS) often depends on fecundity (number of gametes), while male reproductive success is driven by access to mates (Andersson & Iwasa 1996; Bateman 1948), resulting in sex-specific dispersal costs (Hutchings & Gerber 2002; Trochet *et al*. 2016) and highly variable patterns of reproductive variance (Emlen & Oring 1977; Martinig *et al*. 2020; Serbezov *et al*. 2010). Understanding how RS shapes local adaptation is crucial today, as the balance between immigration and selection determines population connectivity and evolutionary potential under ongoing environmental change, providing crucial insights into diversity maintenance, climate responses, and speciation processes (Kawecki & Ebert 2004; Lenormand 2002; Savolainen *et al*. 2007).

Measuring local adaptation in natural environments remains challenging, particularly when comparing the reproductive success of immigrants and local residents. Reciprocal transplant and common garden experiments, though widely used to assess local adaptation, are often impractical for ethical or logistical reasons, especially in protected species (Blanquart *et al*. 2013; Kawecki & Ebert 2004). These challenges are even greater in natural settings, where identifying dispersal events and RS remains particularly difficult.

Field-based pedigrees, which can be genetically informed, offer powerful alternatives to compare fitness between immigrant and local individuals in nature, and have been assessed across diverse taxonomic groups (Bonte *et al*. 2012), revealing consistent differences in birds (Barbraud & Delord 2021; Dickel *et al*. 2024; Hansson *et al*. 2004; Marr *et al*. 2002; Reid *et al*. 2024), mammals (Hasselgren *et al*. 2018; Martinig *et al*. 2020), and fishes (Mobley *et al*. 2019; Peterson *et al*. 2014). Yet, how immigrants*’* fitness varies with their geographic or ecological origin remains poorly understood. Most pedigree-based studies have treated all non-local individuals as a homogeneous group (but see Marr *et al*. 2002 and Peterson *et al*. 2014), assuming that all immigrants experience the same fitness consequences. This approach overlooks potential variation in ecological and genetic properties among source populations that could significantly influence dispersal outcomes and local adaptation patterns.

Furthermore, while the relationship between mating success and RS have been extensively studied to quantify sex-specific selection differences (i.e. Bateman gradient; Andersson & Iwasa 1996; Bateman 1948; Mobley *et al*. 2020; Serbezov *et al*. 2010; Tonnabel *et al*. 2019), studies examining how Bateman gradients vary between immigrant and local individuals remain absent to our knowledge. Indeed, initial dispersal costs experienced by immigrants could potentially be compensated by mating with numerous genetically differentiated individuals (Tallmon *et al*. 2004), maximizing heterosis effects across multiple offspring cohorts while minimizing the risk of a single maladapted pairing.

Anadromous and strongly philopatric species such as the Atlantic salmon (*Salmo salar*) provide ideal models for studying the interplay between dispersal and local adaptation. Atlantic salmon spawn in freshwater, and after 1 to 2 years in freshwater (in France), juveniles migrate to sea for one to three years before returning to their natal river to reproduce (Dumas & Prouzet 2003). This combination of strong philopatry, life-history variation, and environmental heterogeneity among rivers is expected to drive pronounced local adaptation. Yet, substantial proportions of adults can disperse to other rivers, indicating that populations are not fully isolated (Perrier *et al*. 2013). This combination of features results in sufficiently strong population genetic divergence to allow identification not just of immigrating individuals, but also their population of origin, using population genetic individual assignment methods, as well as their RS using genetic parentage testing. This allowed us to quantify origin-dependent fitness differences between immigrants and local individuals in a natural population of Atlantic salmon from the Nivelle River, France. Combining genetic population assignment and pedigree reconstruction over 17 cohort years (>1,100 adults, >3,400 juveniles), we identified immigrants within the population, and investigated the RS of immigrants vs local individuals in relation to sex, body size and sea age. Specifically, we first tested for sex-specific dispersal patterns in relation to sea age, predicting male-biased dispersal as predicted by Local Mate Competition (LMC) theory (Hamilton 1967; Hutchings & Gerber 2002; Trochet *et al*. 2016), earlier maturity of males than females (Barson *et al*. 2015; Mobley *et al*. 2020) and consequently lower sea age at maturity among immigrants. Second, we compared RS of local and immigrant fish, anticipating reduced fitness in immigrants (Fraser *et al*. 2011; Mobley *et al*. 2019; Peterson *et al*. 2014) with variations among source populations. Third, we assessed the fitness of various cross types (local × local, local × immigrant, immigrant × immigrant), expecting heterosis in mixed crosses (Tallmon *et al*. 2004). Finally, we examined Bateman gradients to determine how mating success relates to RS across dispersal phenotypes, predicting steeper Bateman gradients in immigrants (Evans *et al*. 2011; Philippi & Seger 1989). We discuss how origin, sex, and dispersal may interact to shape reproductive success, highlighting the interplay between local adaptation, sexual selection, and gene flow in natural populations.

## Materials and Methods

### Study site

The Nivelle River (39 km, mean flow 4.7 m^3^ s^−1^) supports a self-sustaining Atlantic salmon population at the southern limit of the species range (Fig. 1). Two neighbouring catchments that also support Atlantic salmon populations lie within 20 km of the Nivelle River: the Bidasoa River (69 km long), located about 10 km to the south, and the Adour basin, about 20 km to the north. The latter is subdivided into the Nive and Gave rivers. Other coastal rivers lie >100 km away and were excluded as potential sources of dispersers in this study given typical dispersal distances occur within 10–50 km (Keefer & Caudill, 2014).

**Figure 1.**
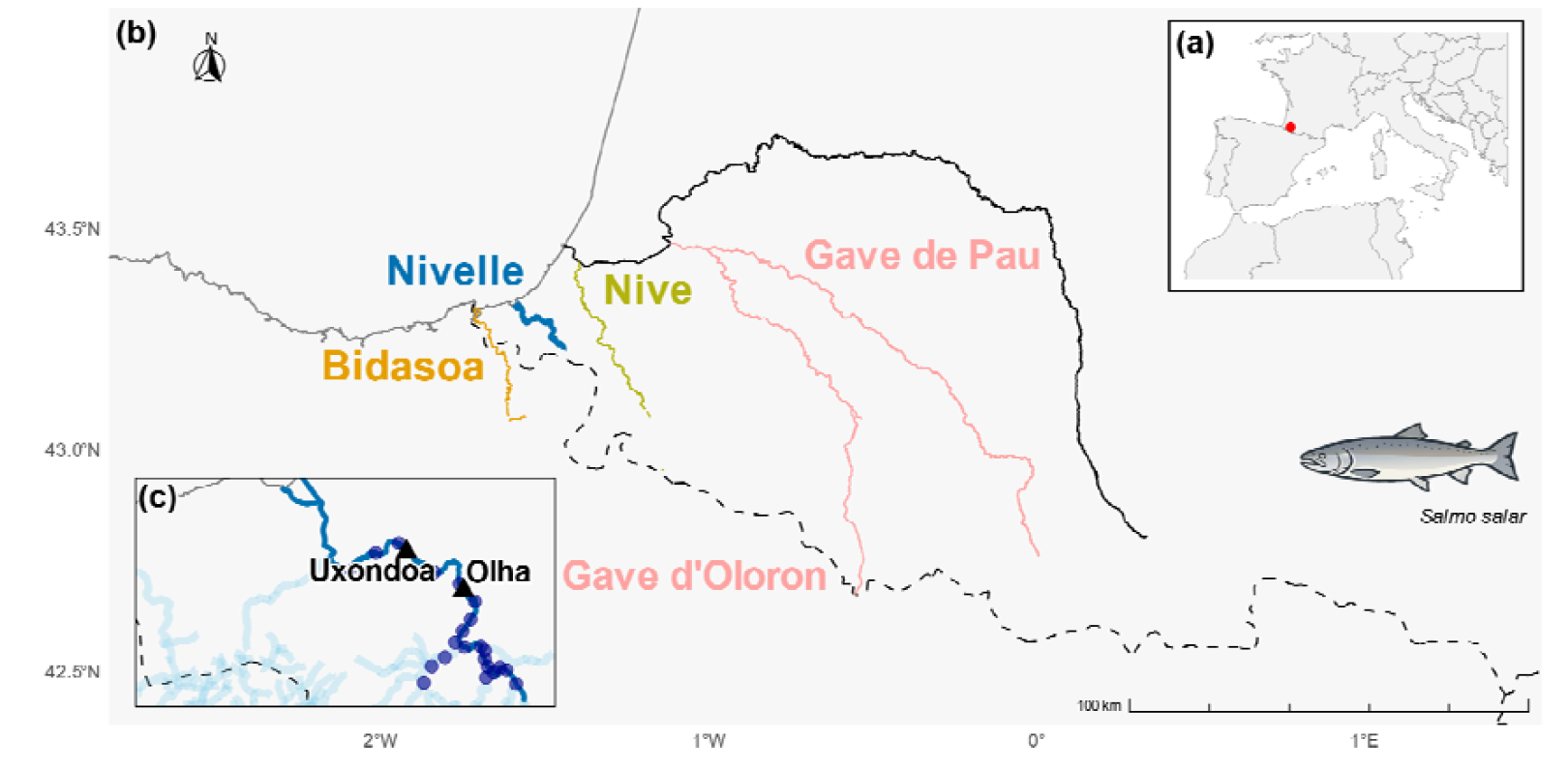
Map of sampling locations for returning adult *Salmo salar* and juveniles in the River Nivelle, southern France. (A) Geographic context with a zoomed inset locating the study area (⍰). (B) Regional overview showing the Nivelle River (blue) and adjacent rivers: Bidasoa (yellow), Nive (green), Gave de Pau and Gave d*’*Oloron (pink). The solid black line represents the Adour River, and the black-dotted line marks the France-Spain border. (C) Detailed map of the Nivelle River displaying sampling sites: ⍰ marks impassable dams at Uxondoa and Olha, both equipped with fish traps for year-round monitoring of returning adults; ⍰ indicates electrofishing sites for juveniles. Deep blue lines represent the main river and its major tributaries, while light blue lines show smaller tributaries in the area.

### Stocking programmes

Between 1972 and 1988, Nivelle was stocked with farmed fry and smolts from Scottish and then Nivelle broodstock (268700 individuals; Baglinière *et al*. 1990); the population is now self-sustaining. Supportive breeding continues in Bidasoa and the Gave de Pau, where adipose-clipped yearlings from local wild progenitors are released. Despite a low return rate, adults of hatchery origin accounted for, on average, one-third of the annual returns to the Bidasoa River between 1993 and 2023 (García-Vega *et al*. 2025).

### Sampling

Since 1991, returning adults in the Nivelle River have been captured at two traps (Fig. 1). Each fish was anaesthetised, measured for fork length (cm), weighed (to the nearest gram), tagged or checked for an existing tag, and fin-clipped for genetic analysis. Adipose fin-clipped individuals were recorded as hatchery fish. Scales were collected from all adults to determine their age in both freshwater and marine environments. Fish that spend one year at sea before returning to the river are classified as one-sea-winter (1SW) individuals, whereas those remaining at sea for more than one year are designated as multi-sea-winter (MSW) individuals. We determined each fish*’*s sex using a molecular method targeting a Y-chromosome-linked marker specific to salmonids (Yano *et al*. 2013, see Supplementary Materials). Juveniles were sampled by electrofishing along the Nivelle River (22 sampling sites) every year in September, and fin-clipped for genetic analysis. Additionally, juveniles from adjacent rivers (Nive, Gave, and Bidasoa rivers) were sampled in 2021 using the same protocol, in order to include them in the baseline for population genetic assignment analyses.

### Molecular analyses

We genotyped a total of 1,175 returning adults including 65 hatchery-born individuals identified by adipose fin clipping, along with 3,281 juveniles captured in the Nivelle River network and 192 juveniles from adjacent rivers. We extracted DNA from fin clips or scales using standard protocols (see Supplementary Materials). We genotyped individuals using either a 176-SNP panel (2014–2019, Aykanat *et al*. 2016) and/or 14 microsatellites (2003-2013, Bacles *et al*. 2018), and retained 4,331 individuals at 164 SNPs and/or 14 microsatellites after quality filtering (Table S1).

### Parentage assignment

We reconstructed pedigrees using FRANz v2.0.0 with a Bayesian sibship method that accounts for unsampled parents (Riester *et al*. 2009). We combined SNP and microsatellite genotypes, set female reproductive age range to 3–6 years and male to 1–6 years, to account for the potential contribution of mature male juveniles to reproduction, a phenomenon known to be significant in some Atlantic salmon populations (Perrier *et al*. 2014). We specified the maximum candidate parents per sex based on abundance estimates (Buoro *et al*. 2019, see Supplementary Materials). We applied a conservative genotyping error rate of 1%, following the recommendations of Janowitz-Koch *et al*. 2019 and Shedd *et al*. 2022, despite observed error rates being lower (<0.1%), and a minimum of 10 shared loci per comparison. Pedigree reconstruction used a Markov Chain Monte Carlo (MCMC) approach with 500 000 burn-in and 3 000 000 sampling iterations; assignments <80% posterior probability or >3 genotype mismatches were treated as missing. Adults and juveniles were then classified by the number of assigned parents (0, 1, or 2). The estimated exclusion probabilities were all above 0.999 for both single-parent and parent-pair exclusions, confirming the high resolution of the marker panel.

### Identifications of immigrants

We assigned adults to one of four source populations (Nivelle, Bidasoa, Nive, Gaves) using assignPOP v1.3.0 (Chen *et al*. 2018), balancing baseline sample sizes by randomly subsampling up to 50 juveniles per source to minimize potential biases in hierarchical structure (Puechmaille 2016). We assessed baseline accuracy via Monte Carlo cross-validation (9 000 iterations) and K-fold validation (K=3–5). Baseline individuals with highest membership probabilities were retained to build final models with a Naive Bayes classification method and predict the origin of 1058 adults. Adults with posterior probability q ≥ 0.75 were classified by source; hatchery fish were classified directly in the *“*disperser*”* category, regardless of identification analyses. We then cross-checked assignments with pedigree data (Table 1; Fig. S5) to determine dispersal status (local vs. immigrant). Immigrants proportion was calculated as the proportion of adults for each population assigned away from Nivelle by the total adult sample size.

**Table 1.** Origin-classification of 722 individuals using an integrative approach combining population assignment tests and pedigree information. Each adult individual was assigned to a predicted source population based on membership probability (*q*), with assignments considered reliable for *q*>0.75. Individuals were further classified by life stage at sampling (*“*A*”* for adult) and by the number of assigned parents: 0A, 1A, 2A for adults (unrelated, one parent, two parents). Only adults assigned to 0 parents and a non-local population (q > 0.75) were considered *‘*true*’* immigrants; those with at least one assigned parent but a genetic background different from Nivelle were excluded from classification to enhance robustness (*“*X*”* individuals). For example, adults with q > 0.75 for Nivelle and at least one assigned parent were considered *‘*true*’* locals. *“*Unrelated-Local*”* denotes individuals with q > 0.75 for Nivelle and no assigned parent. *“*Immigrant F1*”* refers to individuals with at least one immigrant parent and with q > 0.75 for a source different from Nivelle.

| Predicted source population ($q > 0.75$) | | | | |
| --- | --- | --- | --- | --- |
| Parental allocation | Nivelle | Bidasoa | Gave | Nive |
| 0A | Unrelated-local | Immigrant-Bidasoa | Immigrant-Gave | Immigrant-Nive |
| 1A | Local | X | X | X |
| 2A | Local | Immigrant-F1 | Immigrant-F1 | Immigrant-F1 |

### Phenotypic differences between immigrant and local individuals

We tested whether immigrants*’* proportion differed according to sex, sea age, and freshwater age using separate binomial Generalized Linear Models (GLMs; glm, R Core Team, 2023) with origin as response and each trait as predictor. Sea-age categories (1SW, MSW) and sex × sea-age interactions were modelled separately. Pairwise differences in sea-age proportions were evaluated with Bonferroni-corrected post-hoc tests.

### Genetic structure

We estimated genetic differentiation (pairwise F_ST_) among 722 adults confidently assigned to source populations using the θ estimator of Weir & Cockerham 1984 implemented in the R package StAMPP (Pembleton *et al*. 2013) with 10 000 bootstrap replicates to derive 95% confidence intervals (CIs).

### Determinants of reproductive success

For adults sampled from 2004 to 2018, we modeled the relationship between parental traits and reproductive success (RS, offspring count) using zero-inflated Poisson GLMs (pscl, Jackman 2024; Zeileis *et al*. 2008) to account for excess zeros. The model included a binary component for reproduction probability (binomial, logit link) and a count component for offspring number (Poisson, log link). Predictor variables (sex, sea age, body size, origin) were tested in separate models to avoid collinearity between correlated predictors with log(number of offspring sampled) as offset in both zero-inflation and count components. We conducted post-hoc origin effect comparisons using Tukey adjustment (emmeans v 1.10.7, Russell V. *et al*. 2025). We also examined sex-specific and sea-age-specific origin effects to determine whether reproductive fitness differences between immigrants and locals varied across origin categories. Additionally, we fitted a 1SW-only model to isolate local adaptation effects from the inherently higher fecundity of multi-sea-winter (MSW) fish. Among successful breeders (≥1 offspring), we fitted Poisson GLMs (R glm, R Core Team, 2023) by sex to test whether origin effects on offspring number persisted independently of breeding probability. Body-mass differences among origin categories were assessed with Welch*’*s tests.

### Mating patterns

We analysed mating success (unique mates) and Bateman gradients (offspring vs. mates) using zero-inflated GLMs (glmmTMB, Brooks *et al*. 2017) with origin, sea age, and sex as predictors (separate and combined models), including log(number of offspring sampled) as offsets. We defined mating success as the number of unique mates identified for each parent based on pedigree reconstruction. In cases where we identified only one parent, we assumed a single mate. We recorded the type of cross (local x local, mixed, or immigrant x immigrant) and the resulting RS for 141 pairs in which both parents were present. For each pair, we noted the sex, sea age, number of F1 offspring, and origin of each individual. We evaluated if the RS differed among cross types using a negative binomial regression (glmmTMB, Brooks *et al*. 2017) to deal with overdispersion.

The estimated effects from GLMs were expressed as odds ratios (OR) for logistic regressions, calculated by exponentiating model coefficients. For zero-inflated and count models, we exponentiated Poisson coefficients to obtain Rate Ratio (RR), equivalent to a Relative RS. We used R 4.3.1 for analyses and to produce figures (R Development Core Team 2023). We estimated confidence intervals (95% CI) and p-values for all parameter estimates. We set significance levels at α = 0.05.

## RESULTS

### Parentage assignment

Parentage assignment was higher in juveniles than adults, influenced by genotyping error rates and sibship inclusion (Table S2). With a 1% error rate, we generated 197,344 pedigrees (11.9% acceptance). Based on pedigree reconstruction, virtually all females and approximately 80% of males in the true population were represented in our sample (estimated sampling rates: 100% and 80.1%, respectively), with 15.4% unrelated juveniles and 46.9% unrelated adults.

### Population assignment and immigrants identification

Population assignment accuracy increased with training dataset proportion: 91.9% (50% training) to 97.2% (90% training), with population-specific accuracies ranging 90.2-98.1% (Figs. S1-S4). Among 1,058 adults, 768 were confidently assigned (*q*>0.75), including 478 to the Nivelle River (45.2%). After pedigree integration (exclusion of adults with one or both parents assigned), the total immigrants*’* proportion was estimated to be 19.5%: 5.1% (n = 54 individuals) from the Gave source population, 4.2% (n = 45) from the Nive, and 10.2% (n = 108) from the Bidasoa (Table S3 and Fig. S5). All 65 hatchery fish were assigned to Nivelle (52.3%), Bidasoa (24.6%), Nive (16.9%) and Gaves (6.2%). Annual dispersal proportion varied from 21.2% (2012) to 50.0% (2019; Fig. S6). F_ST_ analysis revealed locals genetically closest to unrelated-locals (0.007 ± 0.001) and immigrant-F1s (0.018 ± 0.006), with Nive immigrants most differentiated (0.038 ± 0.01) (Fig. S7).

### Immigrant characteristics

The proportion of immigrants was lower in the MSW category compared to the 1SW category (23.6% vs 37.2%; Odds Ratio = 0.59, 95% CI: 0.40–0.85, p = 0.006). The proportion of immigrants was higher in males than females (39.1% vs 28.1%; OR = 1.57, 95% CI: 1.15–2.16, p = 0.005; Fig. 2). When models were split into sex and sea age categories, a significant higher proportion was found only for the MSW category (41.4% vs 20.7%; OR = 2.67, 95% CI: 1.14–6.12, p = 0.021) despite comprising only 8.4% of males (29 out of 345 males). The proportion of immigrants did not change with freshwater age (Table S4.2).

**Figure 2.**
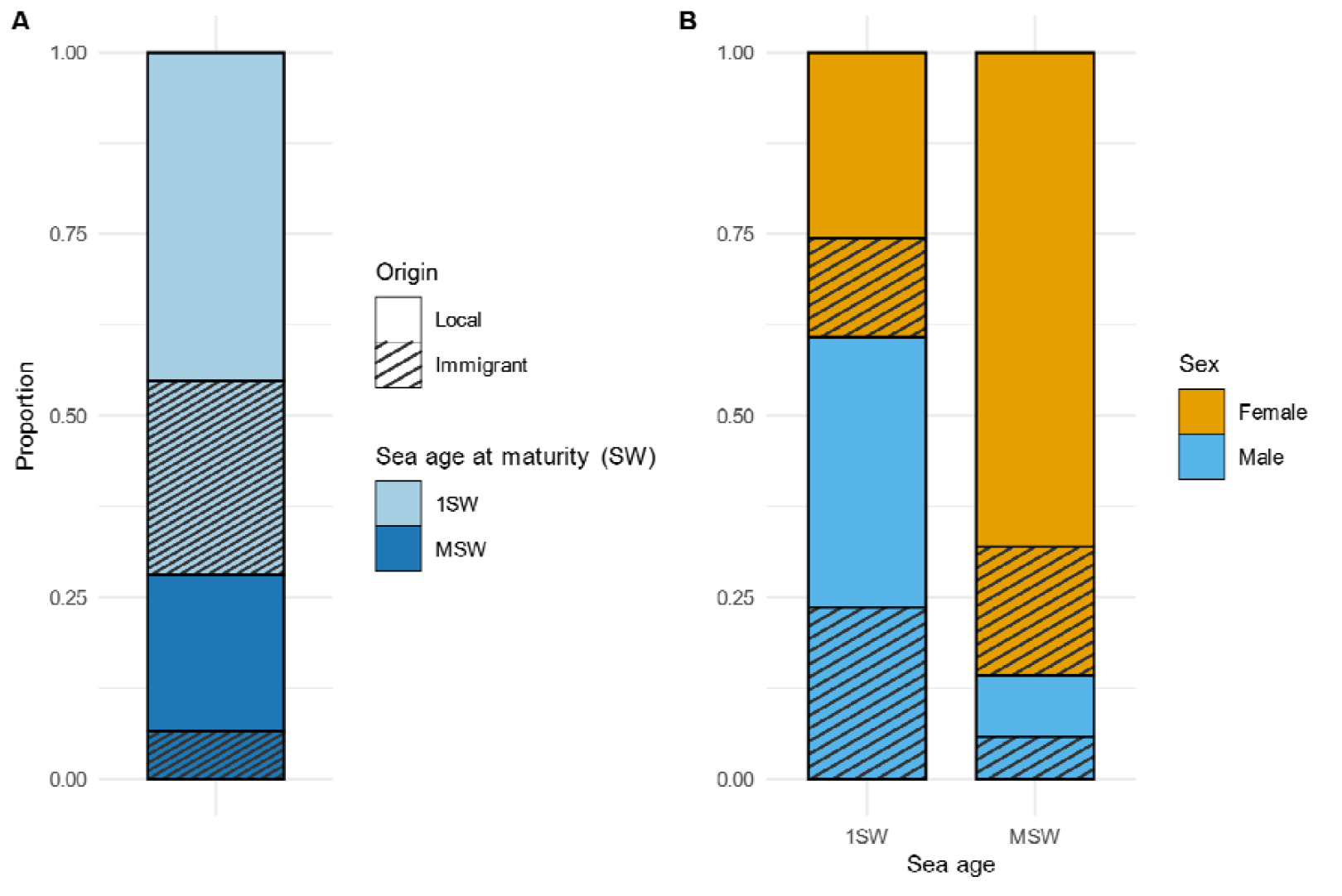
Phenotypic composition of 722 adult salmon confidently classified by origin (Local or Immigrant). (A) Stacked bar plot shows the proportion of 1-sea-winter (1SW, light blue) and multi-sea-winter (MSW, dark blue) adults in Local and Immigrant groups. Local individuals refer to wild-born adults assigned to the Nivelle with population assignment tests and with at least one assigned parent in the pedigree reconstruction; Immigrants represented by hatched shading correspond to genetically assigned dispersers from outside the population and without assigned parents. (B) Proportions of males (blue) and females (orange) within 1SW and MSW groups.

### Reproductive success

Overall, immigrants showed 15% lower RS than locals (Fig. 3A; RR = 0.85, 95% CI: 0.78–0.93, P < 0.001) but this result was mainly driven by male immigrants (RR = 0.70, 95% CI: 0.61–0.81, P < 0.001), whereas immigrant females didn*’*t exhibit significant differences (RR = 1.05, 95% CI: 0.94–1.17, P = 0.40). For origin-specific patterns (Fig.4), female immigrants matched local performance except for Nive (RR = 0.67, 95% CI: 0.47–0.94, *P* = 0.021) and immigrant-F1 (RR = 0.55, 95% CI: 0.44–0.7, P < 0.001). Male immigrants showed lower RS than locals except for Gave, with Nive showing the greatest deficit (RR = 0.28, 95% CI: 0.17–0.47, *P* < 0.001). Among 1SW fish, hatchery origin females showed higher RS than locals (RR = 1.70, 95% CI: 1.32–2.20, *P* < 0.001) with higher mass (2852 ± 418 g vs. 2001 ± 74 g; Tukey HSD, p = 0.029), while three male immigrant categories showed lower performances (Table S7.3). MSW Bidasoa females outperformed locals (RR = 1.76, 95% CI: 1.45–2.15, *P* < 0.001) without significant mass differences (4662 vs. 4395 g, p = 0.229). Across sea age classes, Nive showed consistent disadvantages while hatchery and Gave origins were similar to locals (Table S7.5). MSW Bidasoa fish exceeded local performance (RR = 1.38, 95% CI: 1.14–1.69, *P* = 0.001) without mass differences (4464 vs 4433 g, p= 0.897). An increase of 100% in body mass doubled the reproductive success (Rate ratio = 2.00, 95% CI: 1.83–2.18, *p* < 0.001). This positive relationship between body mass and RS remained significant when analyses were restricted to 1SW individuals (RR = 2.09, 95% CI: 1.76–2.48, *p* < 0.001). Males exhibited 27% lower RS than females (RR = 0.73, 95% CI: 0.67–0.79, p < 0.001). As a consequence, MSW individuals (generally larger because they spent multiple years at sea and predominantly female) exhibited 62% higher RS (RR = 1.62, 95% CI: 1.51– 1.75, *p* < 0.001) compared to 1SW individuals (generally smaller because they spent only one year at sea). Sea-age interactions revealed 18% reduction in 1SW immigrants RS (RR = 0.82, 95% CI: 0.74– 0.92, P < 0.001) but 22% increase in MSW immigrants (RR = 1.22, 95% CI: 1.06–1.40, P = 0.007).

**Figure 3.**
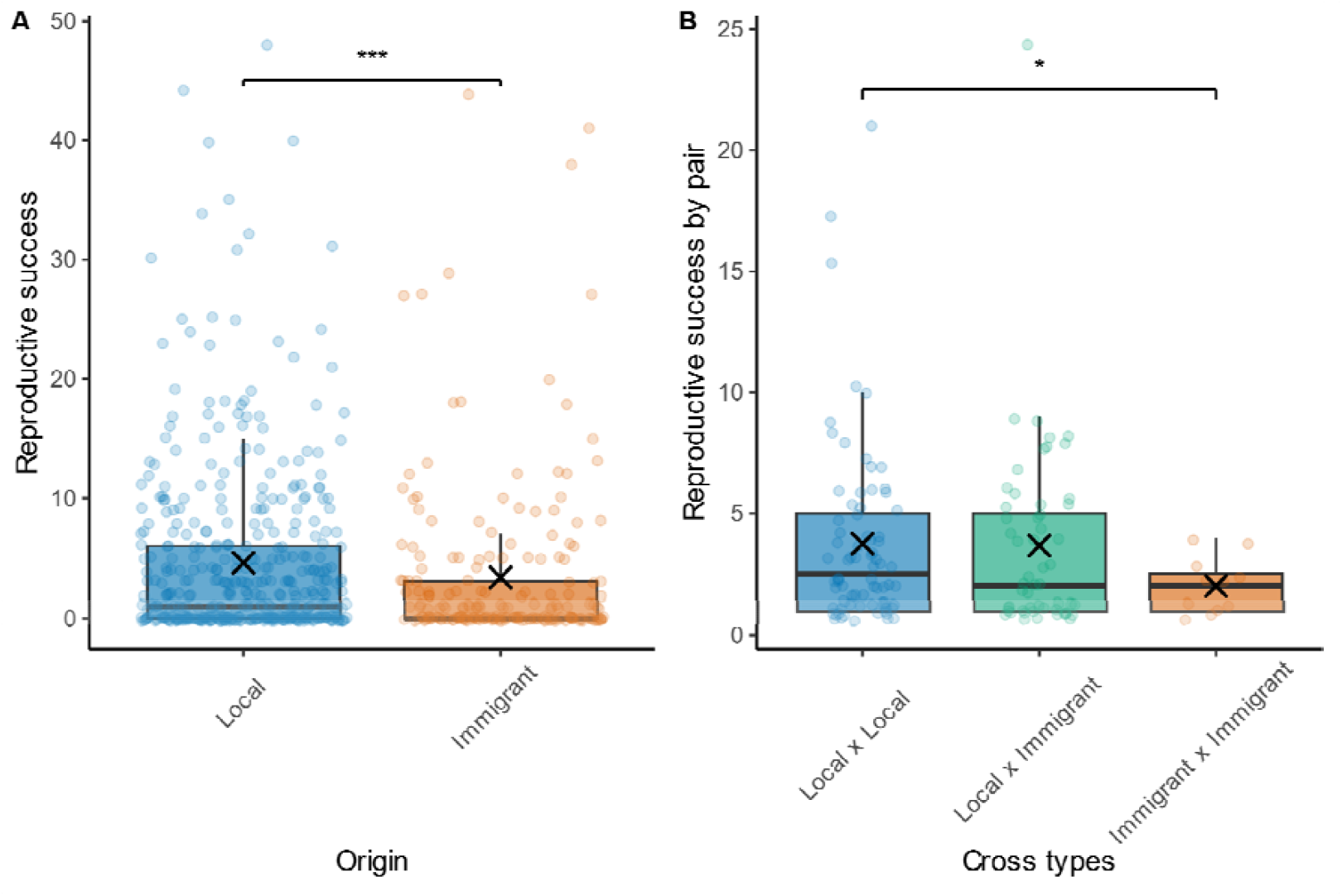
(A) Relationship between adult origin (Local or Immigrant) and individual reproductive success (measured as number of offspring assigned) for 658 adult salmon, (B) Reproductive success per pair for different cross types: Local × Local, Local × Immigrant, and Immigrant × Immigrant. Large X symbols represent means; small circles indicate individual data points. Data points are jittered along the x-axis for clarity. Asterisks indicate statistically significant differences: * * *p < 0.001 and *p < 0.05.

### Mating patterns

Mating patterns varied notably according to origin, sea age, and pair composition (Table S8). Immigrants consistently displayed reduced mating success, securing on average 18% fewer mates than locals (RR = 0.82, 95% CI: 0.69–0.97, p = 0.020). Among sea age classes, MSW individuals obtained 52% more mates than 1SW counterparts (RR = 1.52, 95% CI: 1.30–1.76, p < 0.001). For origin-specific patterns, locals outperformed immigrants for mating success consistently across sexes and sea ages (Table S8.4-5). Analysis of cross types revealed that mixed pairs (N = 54) matched the reproductive output of local pairs (N = 76; RR = 0.98, 95% CI: 0.73–1.31, *p* = 0.90), while immigrant pairs produced 47% fewer offspring (RR = 0.53, 95% CI: 0.29–0.97, *p* = 0.041; Fig. 3B). Sexual selection patterns, as captured by Bateman gradients, showed steeper slopes for males (68% vs 42% per mate; RR = 1.42, 95% CI [1.38, 1.46], p < 0.001; 1.42 × 1.18 = 1.68; Fig. S10 and Table S10.1). Origin-specific analysis revealed immigrant males had steepest gradients with a 94% increase per mate (1.40 × 1.10 × 1.06 × 1.19 = 1.94, p = 0.001; Fig. 5 and Table S10.2) despite 53% lower baseline success (RR = 0.47 for sex [M]×origin [immigrant], 95% CI [0.31, 0.70], p < 0.001). By contrast, local and immigrant females showed intermediate slopes (40 vs. 48% per mate; Table S10.2).

## DISCUSSION

Our study shows that, despite a high proportion of immigrants in this Atlantic salmon population, their contribution to gene flow is not uniform but depends strongly on origin, sex, and life-history traits. While male immigrants consistently showed reduced reproductive success, female immigrants often matched or exceeded local performance, particularly among MSW individuals. These results highlight sex-specific selective pressures, with immigrant males experiencing stronger sexual selection intensity through steeper Bateman gradients and increased variance in mating success, consistent with local mate competition theory (Li & Kokko 2019). Origin effects were striking: Nive immigrants suffered severe reproductive disadvantages, while Gave and hatchery fish performed comparably to locals, and Bidasoa females even outperformed residents. Reduced fitness of immigrant × immigrant crosses further underscores the buffering role of local genotypes in maintaining adaptive potential. Together, our findings reveal complex interactions between dispersal, sexual selection, and local adaptation, showing immigrant contributions to gene flow and evolutionary dynamics are more heterogeneous than often assumed.

### Sex-biased dispersal

By combining population assignment and parentage analysis, we successfully identified immigrants and their population of origin, despite substantial gene flow. We detected a high overall immigration rate (19.5%), varying from 4.2% (Nive) to 10.2% (Bidasoa). These results differ from earlier findings, based on a different methodology, reporting that 12–23% of Nivelle salmon originated from the Bidasoa between 1990 and 1996, and none from the Nive (Valiente *et al*. 2010). Geographic distance may explain these different dispersal rates among source populations but demography may also matter: higher adult returns reduce dispersal (Hard & Heard 1999; Quinn & Fresh 1984; Westley *et al*. 2025), while dispersal tends to decrease with larger basins (Chat *et al*. 2022; Westley *et al*. 2025). Interannual variation in immigration (Fig. S6) highlights the interplay of genetic predisposition, environment, and demography in shaping dispersal. A caveat is that some detected immigrants may not be true spawners but instead exploratory individuals assessing spawning habitats (Ahnesjö & Forsman 2006; Keefer & Caudill 2014).

We documented male-biased immigration (Fig. 2), consistent with some salmonid studies (Bekkevold *et al*. 2004; Hamann & Kennedy 2012) but contrasting others (Consuegra & Garcia de Leaniz 2008; Mobley *et al*. 2019; Pollock *et al*. 2020), emphasizing its context-dependent nature. Stronger sexual selection in males (Fig. 5 and Fig. S10) supports local mate competition (LMC) theory, which predicts higher dispersal in the sex experiencing greater intrasexual competition (Hamilton 1967; Lawson Handley & Perrin 2007; Li & Kokko 2019). Conversely, the sex securing territories usually faces higher dispersal costs, favouring philopatry near natal sites (Li & Kokko 2019); indeed, salmon males compete for mates, while females invest in redds (Fleming & Gross 1989). However, we found no greater female dispersal costs (Fig. 4A). Theoretical models predict greater dispersal in the sex with lower costs (here, males) (Gros *et al*. 2008). Male biased dispersal may also reflect inbreeding avoidance, since only one sex needs to disperse (Li & Kokko 2019), and females capable of rejecting inbred matings would favour male dispersal to seek unrelated mates (Keller & Waller 2002). The predominance of 1SW immigrants is expected because there were more male immigrants, and males mature earlier than females (Barson *et al*. 2015; Mobley *et al*. 2020). Male bias among MSW fish (Fig. 2) may stem from size-related advantages (Bonte *et al*. 2012; Merckx *et al*. 2018; Roff 1977).

**Figure 4.**
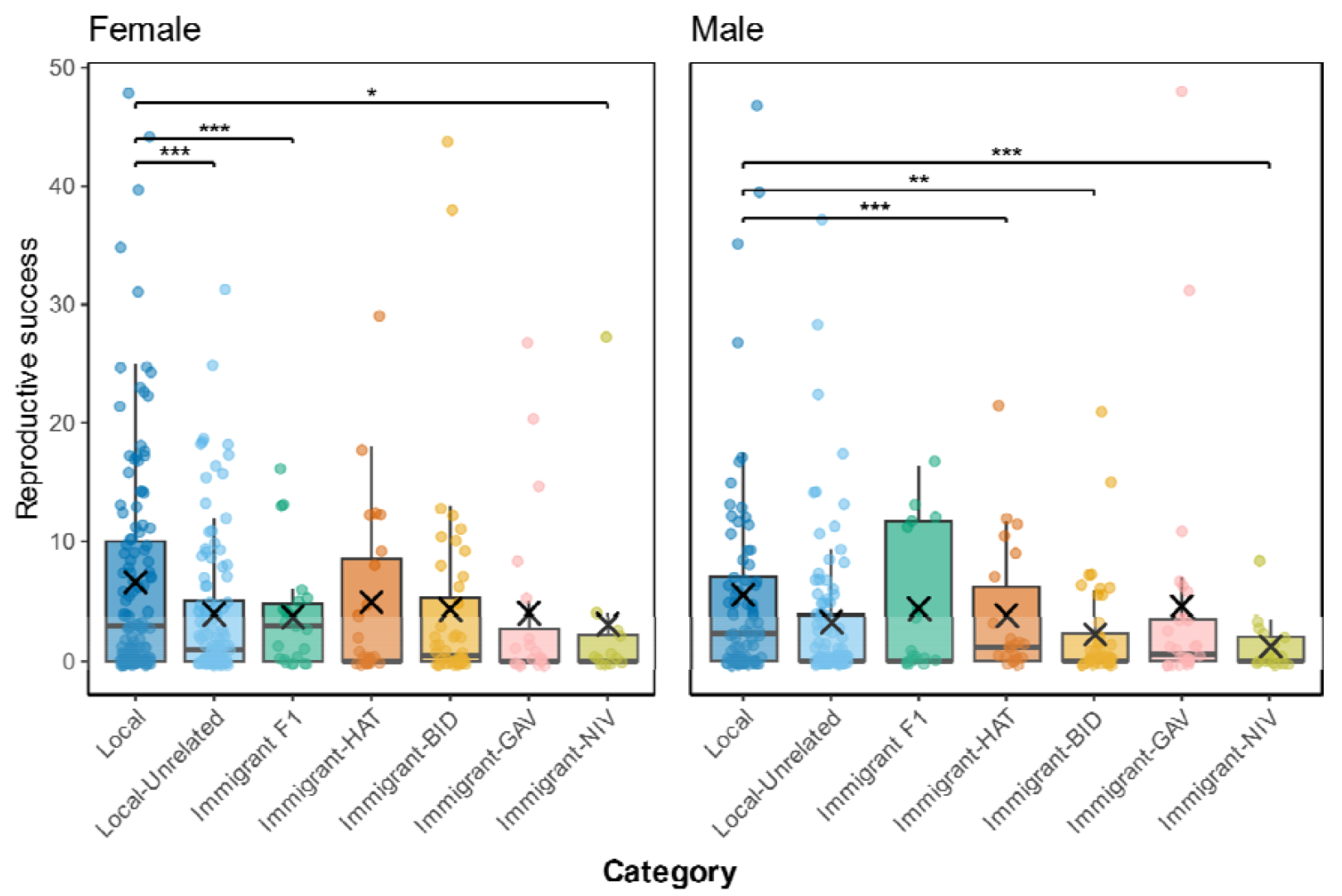
Reproductive success (number of assigned offspring) among female (left, n = 361) and male (right, n = 297) salmon grouped by origin categories. Large *“*X*”* symbols represent group means; small circles show individual data points. Category codes: Local refers to individuals assigned to the Nivelle population (residents); Local-Unrelated corresponds to individuals assigned to Nivelle but without identified parents in pedigree analysis; Immigrant-F1 represents first-generation immigrants with parents assigned in the Nivelle River but genetically assigned to other source populations; HAT, BID, GAV and NIV denote immigrants from Hatchery, Bidasoa, Gave and Nive populations, respectively. Data points are jittered along the x-axis for clarity. Asterisks indicate statistically significant differences: ***p < 0.001, * *p < 0.01, *p < 0.05.

**Figure 5.**
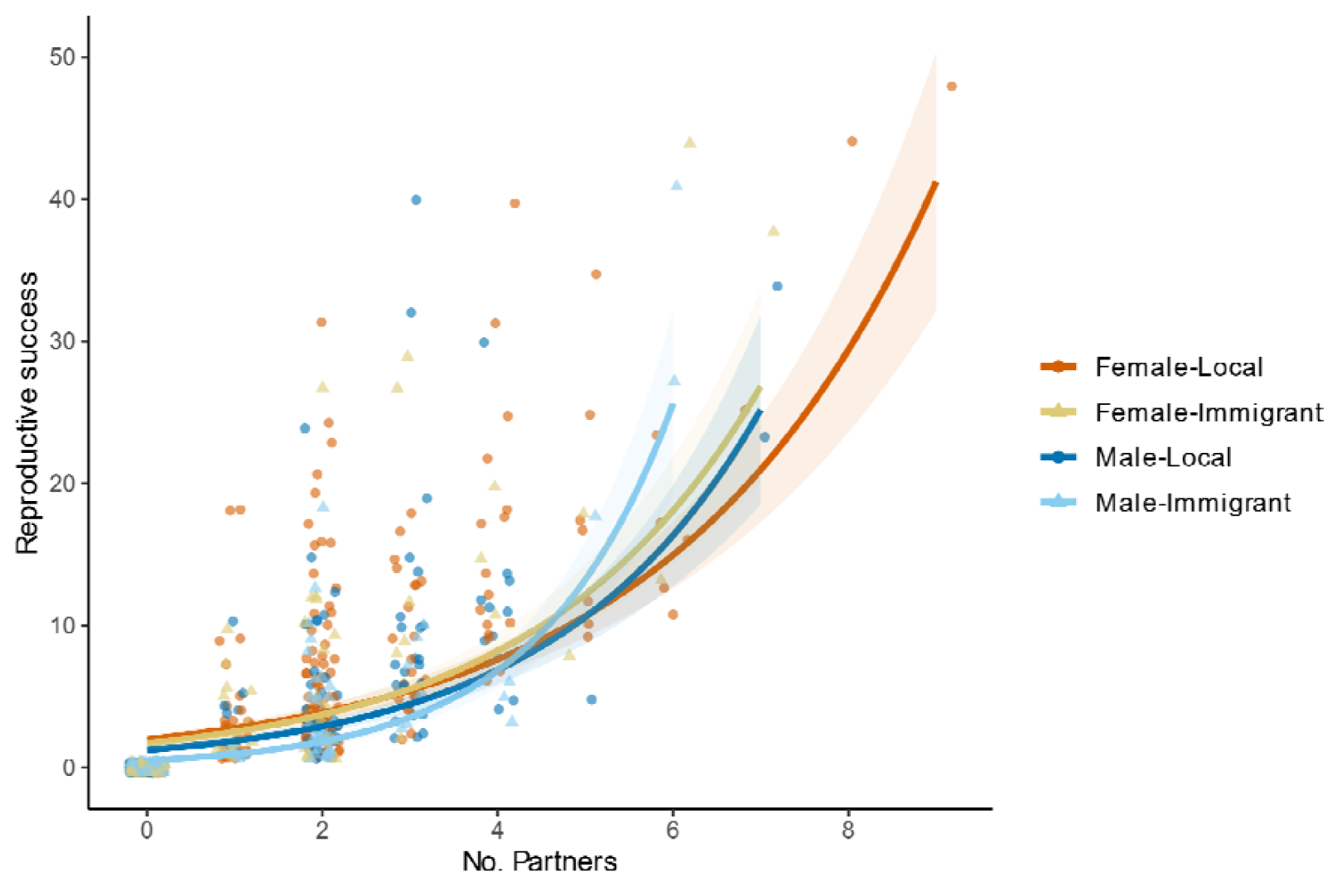
Bateman gradients showing the relationship between reproductive success and number of partners in 658 adult salmon. Points represent individual data for female locals (orange circles), female immigrants (yellow triangles), male locals (dark blue circles), and male immigrants (light blue triangles). Lines and shaded areas show fitted relationships (± SE) for each sex and origin category.

### An origin-dependent fitness

Few studies have compared reproductive success across immigrant origins despite its importance for local adaptation. Immigrants averaged 15% lower RS than locals (Fig. 3A), consistent with selection against dispersers in salmonids (Mobley *et al*. 2019; Peterson *et al*. 2014). RS varied by origin: strong selection against Nive immigrants, intermediate for Bidasoa, and equivalent RS for Gave/Hatchery. Such heterogeneity reflects scale-dependent local adaptation: a salmonid meta-analysis shows stronger adaptation with greater distance, but highly variable outcomes at intermediate scales (~100 ⍰ km) like ours (Fraser *et al*. 2011). RS estimates require caution, as some adults sampled in recipient rivers may have spawned elsewhere, appearing with zero offspring in pedigrees (Keefer & Caudill 2014), but analyses restricted to successful breeders (Fig. S9) confirm our findings. Offspring of immigrant crosses showed reduced RS (Fig. 3B), highlighting the advantage of at least one local genotype (Tallmon *et al*. 2004), though F1 benefits may mask later-generation costs including expression of deleterious recessives and epistatic breakdown (Dickel *et al*. 2024; Hasselgren *et al*. 2024; Marr *et al*. 2002; Martinig *et al*. 2020). If F1 inherit dispersal tendencies, their emigration could further underestimate long-term immigrant contributions (Doligez & Pärt 2008), and complex genetic architectures may amplify this in subsequent generations. The lower RS of most immigrant F1s (Fig. 4A; Fig. S9; Table S7) supports intergenerational fitness costs and underscores the need for long-term pedigree tracking to fully capture evolutionary consequences of immigration (Koch & Narum 2021).

### Sex-specific and life-history dependent reproductive success

Sex and life-history traits shaped fitness outcomes. Males exhibited 27% lower RS than females (Fig. S10; Table S5.4), reflecting intrinsic life-history differences. Females*’* longer marine residence (Barson *et al*. 2015) led to a higher proportion of MSW individuals, which had 62% higher RS than 1SW females, with benefits scaling with age and body mass (Table S5.1; S5.3, Mobley *et al*. 2019, 2020). Notably, immigrant status accentuated the reproductive shortfall in males, potentially reflecting female mate choice or local adaptation processes such as assortative mating (Higashi *et al*. 1999; Snowberg & Benkman 2009; Tobler *et al*. 2009), a component of salmonid behaviour that can affect RS (Auld *et al*. 2019). A study on Atlantic salmon revealed no clear evidence for origin-based assortative mating (Mobley *et al*. 2019), though this examination was constrained by a limited number of dispersers. Contrary to predictions that females—primary territory acquirers and defenders (Fleming & Gross 1989)—should bear the greatest dispersal costs (Li & Kokko 2019; Martinig *et al*. 2020), most female immigrants matched or exceeded local RS (Fig.4A; Table S7), possibly driven by intrinsic factors such as size and condition (Heinimaa & Heinimaa 2003; Koch & Narum 2021; Serbezov *et al*. 2010). MSW immigrant females*’* strong performance despite moderate weight differences underscores the benefit of prolonged marine growth. Unexpectedly, 1SW hatchery fish showed superior RS, likely because local broodstock use minimizes domestication effects (Ford *et al*. 2012; Hess *et al*. 2012), even persisting across generations (Janowitz-Koch *et al*. 2019). However, rapid hatchery introgression can alter key traits such as return timing, with potential for phenotypic mismatch and population-level fitness declines (May *et al*. 2024) and broader dispersal impacts to non-target populations (Lamarins *et al*. 2024; Perrier *et al*. 2013). Our analysis of F1 offspring leaves heterosis persistence in later generations unresolved.

### Differential strength of sexual selection reveals sex-specific reproductive heterogeneity

Males showed higher variance in mating success and offspring number (Fig. 5; Fig. S10, Garant *et al*. 2001; Serbezov *et al*. 2010), consistent with Bateman*’*s principles where female RS primarily depends on offspring production and survival, while male success depends on mate access (Andersson & Iwasa 1996; Bateman 1948). Our system revealed a flexible mating system benefitting both sexes via female polyandry (up to nine mates), reflecting salmonids*’* multiple-redd strategy that spreads spawning effort to increase fertilization opportunities and reduces fitness variance under environmental stochasticity (Barlaup *et al*. 1994; Serbezov *et al*. 2010). Coming from afar could present some advantages in terms of sexual selection. Although immigrant and local females had similar mating success, immigrant males experienced lower baseline RS but faced disproportionate fitness returns when acquiring multiple mates, underscoring sex-dependent dispersal benefits (Li & Kokko 2019). However, the small number of immigrant males with multiple mates limits the robustness of this conclusion. The elevated strength of sexual selection on immigrant males, manifested in their steeper Bateman gradients, may counteract local maladaptation by maximizing beneficial local–immigrant allele combinations, generating disproportionate fitness returns for recipient populations (Tallmon *et al*. 2004).

## CONCLUSION

This study demonstrates that immigrant fitness is origin-dependent, challenging the traditional assumption of uniform dispersal costs. Overlooking this variation risks misestimating the genetic consequences of immigration and misrepresenting local adaptation processes. While locals generally outperform immigrants—reinforcing adaptive differentiation (Nosil *et al*. 2005)—some immigrant groups perform as well as residents, thereby maintaining genetic diversity that might buffer demographic fluctuations and enhance adaptive potential under environmental change. Our results further show that male-biased dispersal occurs despite higher fitness costs, consistent with sex-biased dispersal theory and highlighting the importance of local mate competition. Yet, immigrant males acquiring multiple mates may achieve several reproductive returns, facilitating novel genetic combinations that could prove adaptive in changing environments. These findings underscore the need to better understand the mechanisms driving origin-dependent immigrant fitness and call for quantifying multigenerational effective gene flow—an evolutionarily critical but still understudied process that only long-term pedigree data can fully capture.

## Supporting information

Supplementary Materials for Selection against dispersers: Sex, age and origin-dependent fitness differences in wild Atlantic salmon

## Data availability statement

Raw genotype data and scripts associated used in this study are available in the INRAE data repository

## Acknowledgements

We thank Frédéric Lange, AAPPMA Nivelle, Migradour, Orekan and all those involved in data collection at the Nivelle, Gave, Nive and Bidasoa facilities over the years, and Annukka Ruokolainen for skilled laboratory assistance. We are also grateful to Cecile FE Bacles and Olivier Lepais for providing microsatellite data and to Steven A. Ramm and Kenyon Mobley for helpful discussions on data analyses. The samples used in this study were provided by the Biological Resource Centre Colisa (DOI: Biological Resource Centre Colisa) » part of BRC4Env (DOI: https://doi.org/10.15454/TRBJTB), of the Research Infrastructure AgroBRC-RARe.

