## Supplementary Materials for Selection against dispersers: Sex, age and origin-dependent fitness differences in wild Atlantic salmon for "Selection against dispersers: Sex, age and origin-dependent fitness differences in wild Atlantic salmon"

This PDF file includes:

**Supplemental Text (Page 3)**

**Supplemental Figures (Page 7)**

**Fig S1.** Assignment accuracies estimated via Monte-Carlo cross-validation with a linear discriminant analysis (lda) classification method.

**Fig S2.** Membership probability of the four populations used in the study system.

**Fig S3.** Assignment accuracies estimated after filtering individuals with highest membership probability via Monte-Carlo cross-validation with a linear discriminant analysis (lda) classification method.

**Fig S4.** Membership probability estimated after filtering individuals with the highest membership probability of the four populations used in the study system.

**Fig S5.** Comparison of dispersal rates estimated using population assignment alone versus population assignment combined with pedigree reconstruction.

**Fig S6.** Temporal variation in the proportion of individuals by origin category from 2004 to 2019.

**Fig S7.** Pairwise FST among populations for 722 adults confidently assigned to their population of origin.

**Fig S8.** Sea age proportions (1SW vs MSW) among origin categories for 722 adults confidently assigned to their population of origin.

**Fig S9.** Reproductive success (number of assigned offspring) among female (left, n = 210) and male (right, n = 149) successful breeders salmon grouped by origin categories.

**Fig S10.** Bateman gradients showing the relationship between reproductive success and number of partners in 658 adult salmon.

**Fig S11.** Bateman gradients showing the relationship between reproductive success and number of partners in 359 successful breeders adult salmon.

**Supplemental Tables (Page 18)**

**Table S1.** Number of genotyped individuals by genotyping method, stage of life, and sampling year.

**Table S2.** Summary of parentage assignment runs using FRANz software with different parameter settings.

**Table S3.** Population assignment results using Naïve Bayes classification corrected with pedigree information.

**Table S4.** Summary of generalized linear models (GLMs) testing phenotypic differences between immigrant and local Atlantic salmon adults (n = 722).

**Table S5.** Summary of generalized linear models (GLMs) testing effects of phenotypic traits on reproductive success (RS) in Atlantic salmon adults (n = 658).

**Table S6.** Summary of generalized linear models (GLMs) testing effects of origin on reproductive success (RS) in Atlantic salmon adults (n = 658).

**Table S7.** Summary of generalized linear models (GLMs) testing effects of population origin categories on reproductive success in Atlantic salmon adults (n=658).

**Table S8.** Summary of generalized linear models (GLMs) testing effects of origin, sea age, sex, and origin categories on mating success in Atlantic salmon adults (n=658).

**Table S9.** Summary of generalized linear models (GLMs) testing effects origin and origin categories on reproductive success in successful breeders Atlantic salmon adults (n = 359).

**Table S10.** Summary of zero‐inflated Poisson generalized linear models testing effects of number of partners, sex, origin, and their interactions on reproductive and mating success in Atlantic salmon adults (n=658) and successful breeders (n = 359).

### Supplemental Text

#### *DNA extraction*

We extracted DNA using a salt–chloroform extraction protocol adapted from Müllenbach *et al.* 1989 or Purelink Genomic DNA mini kit (Invitrogen; www.invitrogen.com) following the manufacturer protocol respectively from fin clips and scales. From 2016, we extracted the DNA of scale samples with NucleoSpin 96 Tissue (Macherey-Nagel; [www.mn-net.com](http://www.mn-net.com)) following the manufacturer protocol.

#### *Genotyping*

We carried out different genotyping methods for this study, with some individuals genotyped with a SNP chip, others at microsatellite loci, and some using a combination of the two methods (**Table S1**). We genotyped 463 adults and 1100 juveniles from 2014 to 2019 using 176 Single Nucleotide Polymorphism (SNP) markers developed for Atlantic salmon (Aykanat *et al.* 2016). These markers come from a SNP panel (N = 196 SNPs), designed to estimate the overall genome-wide following level of differentiation among populations within a system (Teno river, Norway). These SNPs included one sex-specific marker linked to the salmonid Y-chromosome (Yano *et al.* 2013) and 5 markers associated with sea age at maturity in *S.salar*. From 2003 to 2013, we genotyped 70 adults and 1238 juveniles using 14 microsatellite markers (Bacles *et al.* 2018). In addition, we also genotyped these individuals with three similar markers to the SNP panel: 1 sex-specific marker, and 2 loci associated with sea age using a PCR-based KASP genotyping assay. We genotyped 642 adults and 943 juveniles with both methods. When we compared the genotypes of the 3 similar markers obtained with SNP panel and KASP assay, we found a 100% match.

We assessed the quality of genotyping by removing individuals and markers with low genotyping success (<80% and <90% for individuals and markers respectively), as well as monomorphic markers. We also identified and excluded the cited previous markers associated with sea age at maturity in *S.salar* using PCADAPT (Luu *et al.* 2017). To ensure accurate sibship reconstruction based solely on genetic data over the 17-years period (during which generations overlap), we included each unique multilocus genotype (MLG) only once in the dataset, thereby excluding recaptures and technical replicates. We verified the discrimination of related individuals using genotyping error estimates and clone analysis, performed with the R package POPPR (Kamvar *et al.* 2014). In total, we successfully genotyped 4331 individuals at 14 microsatellites markers and/or 164 SNP loci and retained them for further analyses.

##### *Microsatellites information*

See Supplementary information of Bacles, C.F.E., Bouchard, C., Lange, F., Manicki, A., Tentelier, C., Lepais, O., 2018. Estimating the effective number of breeders from single parr samples for conservation monitoring of wild populations of Atlantic salmon Salmo salar. J Fish Biol 92, 699–726. <https://doi.org/10.1111/jfb.13537>

##### *SNP array information*

Sex marker (sdy assay) is based on the ratio of sdy reads to the average of the “VIP SNPs” (the 5 listed below). The default thresholds are:

<0.005 = female and >0.02 = male. Anything in between, or when the coverage of any one of the 5 VIP SNPs is <20 is listed as ‘NA’. This may bias the calls towards females.

For all other SNPs, see Supplementary information of Aykanat, T., Lindqvist, M., Pritchard, V.L., Lindqvist, M., Primmer, C.R., 2016. From population genomics to conservation and management: a workflow for targeted analysis of markers identified using genome-wide approaches in Atlantic salmon *Salmo salar*. J. Fish Biol. 89, 2658–2679. <https://doi.org/10.1111/jfb.13149>.

#### *Escapment estimate*

We marked salmon captured at the Uxondoa trap with a passive integrated transponder (PIT-tag) microchip. Recaptures at the Olha trap allowed estimation of trap efficiency and, consequently, provided an index of population size.

Nm_max_ and Nf_max_ define the maximum number of candidate parents (males and females respectively). To define these parameters, we hypothesized that these parameters were approximatively half of the estimates of escapment derived from aerial surveys and stream walks, assuming that approximately half of the fish were male and half were female.

a.	Nm_max_: we take into account the fact that 50% of the parr mature, and can therefore contribute to reproduction (the maximum total number of parr / 2).

b.	Nf_max_: For females, the number of individuals that return to the river to spawn.

#### *Parentage assignment*

FRANz uses a full-probability Bayesian model for pedigree reconstruction, which is particularly advantageous for studies with incomplete sampling of potential parents and offspring. This model accounts for unsampled parents and leverages sibships among sampled individuals to infer parental genotypes from offspring, thereby improving pedigree resolution (Jones et al., 2010; Riester et al., 2009). We tested the effect of combining individual genotypes from our SNP panel and microsatellite data with varying biological information and error rates on parentage assignment (**Table S2**).

Specifically, we set the reproductive age range to 3–6 years for females and 1–6 years for males, to account for the potential contribution of mature male juveniles to reproduction, a phenomenon known to be significant in Southern Atlantic salmon populations (Perrier *et al.* 2014). We also provided the maximum number of candidate parents by sex (N_fmax_ and N_mmax_) using estimates of adults and juvenile abundances in the Nivelle (Buoro et al. 2019). We applied a conservative genotyping error rate of 1%, following the recommendations of Janowitz-Koch et al. (2019) and Shedd et al. (2022), despite observed error rates being lower (<0.1%).  We set the minimum number of common typed loci for a pair of individuals to *10* to take account of different genotyping methods. We conducted pedigree reconstruction using a Markov Chain Monte Carlo (MCMC) approach with 500,000 burn-in iterations followed by 3,000,000 sampling iterations. We considered as missing assigned parents that had a posterior probability of assignment < 80% and mismatches >3. The final parentage assignment enabled classification of individuals based on their life stage (‘A’ for adult, ‘J’ for juvenile) at the time of sampling and the number of parents assigned (either 0 parent: Unrelated (U), or 1 or 2 parents). We classified adults as UA, 1A, or 2A, and “juvenile” as UJ, 1J, or 2J.

#### *Identification of immigrants*

In MCMC approach, we randomly resampled different proportions of individuals from each population (50%, 70%, and 90%) as training sets, with the remaining individuals serving as holdout test sets for each iteration. We used all available markers for each combination of training individuals.

In K-fold cross-validation, individuals from each population were divided into K groups, where one group served as the test set while the remaining K-1 groups formed the training set. This process was repeated until every group had been tested once, ensuring that every individual was evaluated exactly once and received a membership probability estimate. We then extracted the baseline individuals with the highest membership probabilities and re-ran both resampling cross-validation methods using the same associated parameters.

#### *Reproductive success of dispersers*

The zero-inflated was chosen because the Poisson model alone underfit zeros (Observed zeros: 300, Predicted zeros: 56, Ratio: 0.19, p < .001). The negative binomial model displayed an underdispersion (dispersion ratio = 0.551, p-value = 0.04)

#### *Reproductive success among immigrant categories*

When examining reproductive success across population origin categories, local individuals with assigned parents exhibited higher reproductive success than all other categories (Table S7.1), including unrelated locals, F1 immigrants and immigrants from different source populations. Individuals from Bidasoa and Nive showed particularly low reproductive success compared to natives (RR = 0.79, 95% CI: 0.69–0.90, P < 0.001; and RR = 0.47, 95% CI: 0.35–0.63, P = 0.001, respectively), as well as compared to immigrant F1 (RR = 0.71, 95% CI: 0.59–0.84, *P* = 0.001). Contrary to our expectations, there was no significant difference between hatchery-origin immigrants and natives, nor between immigrants from Gave and natives (RR = 0.93, 95% CI: 0.79–1.09, *P* = 0.4; and RR = 0.99, 95% CI: 0.85–1.16, *P* > 0.9).

#### *Pairwise comparisons among immigrant categories*

Nive immigrants consistently demonstrated the poorest reproductive performance, with significant disadvantages across all comparisons (P < 0.0001 for most categories, P = 0.03 for F1 immigrants). In contrast, local individuals achieved superior reproductive success, significantly outperforming F1 immigrants (P = 0.0004), Bidasoa immigrants (P = 0.0005), and unrelated locals (P = 0.0053). Notably, locals performed equivalently to hatchery (P = 0.5253) and Gave immigrants (P = 0.9126), suggesting these immigrant origins may not undergo significant fitness costs.

Some clusters emerged among immigrant categories. F1, hatchery, and Bidasoa immigrants formed a statistically indistinguishable group (all P > 0.05), while hatchery and Gave immigrants showed equivalent performance (P = 0.9990), as did Bidasoa and Gave immigrants (P = 0.5349).

#### *Reproductive success of immigrants among breeders*

Among successful breeders only (210 females, 149 males), immigrants overall showed reduced reproductive success compared to locals (RR = 0.86, 95% CI: 0.79–0.93, P < 0.001). However, this general pattern masked considerable sex- and origin-specific variation. Among females, immigrant F1s showed the strongest reduction in reproductive success (RR = 0.56, 95% CI: 0.44–0.70, P < 0.001), followed by local-unrelated individuals (RR = 0.82, 95% CI: 0.73–0.92, P < 0.001) and Nive immigrants (RR = 0.69, 95% CI: 0.49–0.95, P = 0.025). Other immigrant categories showed no significant difference from locals: hatchery (RR = 1.16, 95% CI: 0.95–1.41, P = 0.15), Bidasoa (RR = 0.90, 95% CI: 0.77–1.06, P = 0.2), and Gave (RR = 0.98, 95% CI: 0.78–1.23, P = 0.8) females all matched local performance. Among males, most immigrant categories remained disadvantaged: Nive showed the strongest reduction (RR = 0.32, 95% CI: 0.20–0.51, P < 0.001), followed by Bidasoa (RR = 0.62, 95% CI: 0.48–0.78, P < 0.001) and hatchery (RR = 0.71, 95% CI: 0.55–0.91, P = 0.007). Only Gave males matched locals (RR = 1.01, 95% CI: 0.81–1.25, P > 0.9), while immigrant F1 males showed no significant difference (RR = 1.11, 95% CI: 0.85–1.45, P = 0.5). These results indicate that the overall 14% reduction in immigrant reproductive success arises from sex- and origin-specific effects rather than a universal local advantage.

#### *Sex-specific Bateman gradients with individual origin among breeders*

The Bateman gradients varied significantly among sex-origin combinations. Incorporating origin alongside sex revealed substantial group-specific differentiation in mating success returns: local females showed a 33% increase per additional mate (RR = 1.33, 95% CI [1.30, 1.36], p < 0.001), local males gained 35% per mate (1.33 × 1.04 = 1.38; RR = 1.04 for partners×sex [M], 95% CI [0.98, 1.10], p = 0.2), immigrant females exhibited a 40% increase per mate (1.33 × 1.05 = 1.40; marginal origin effect RR = 1.05, p = 0.063), and immigrant males displayed the steepest gradient at 56% per additional partner (1.33 × 1.04 × 1.05 × 1.13 = 1.56; RR = 1.13 for partners×sex [M]×origin [immigrant], 95% CI [1.02, 1.25], p = 0.019), despite exhibiting 52% lower baseline reproductive success (RR = 0.48 for sex [M]×origin [immigrant], 95% CI [0.33, 0.70], p < 0.001).

### Supplemental Figures

**Fig S1.** **Assignment accuracies estimated via Monte-Carlo cross-validation with a linear discriminant analysis (lda) classification method.** We used three levels of training individuals (50%, 70% and 90% of individuals from each population, on x-axis). ‘Overall’ represents the whole dataset, ‘Nivelle_P’, JUNIV, JUGAV, and JUBID the subset of juveniles from Nivelle, Nive, Gave and Bidasoa populations respectively. We used all available markers for each combination of training individuals.
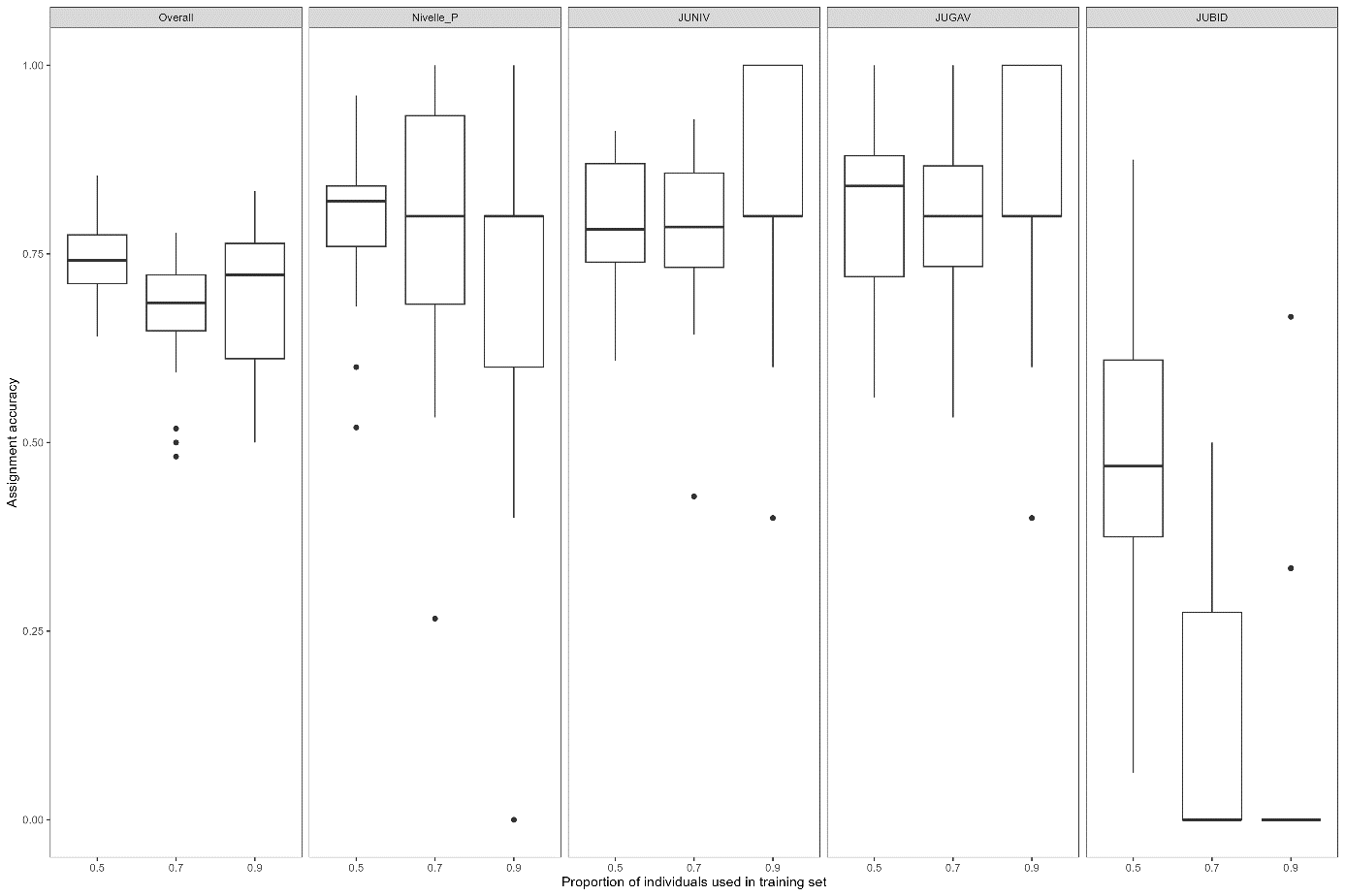

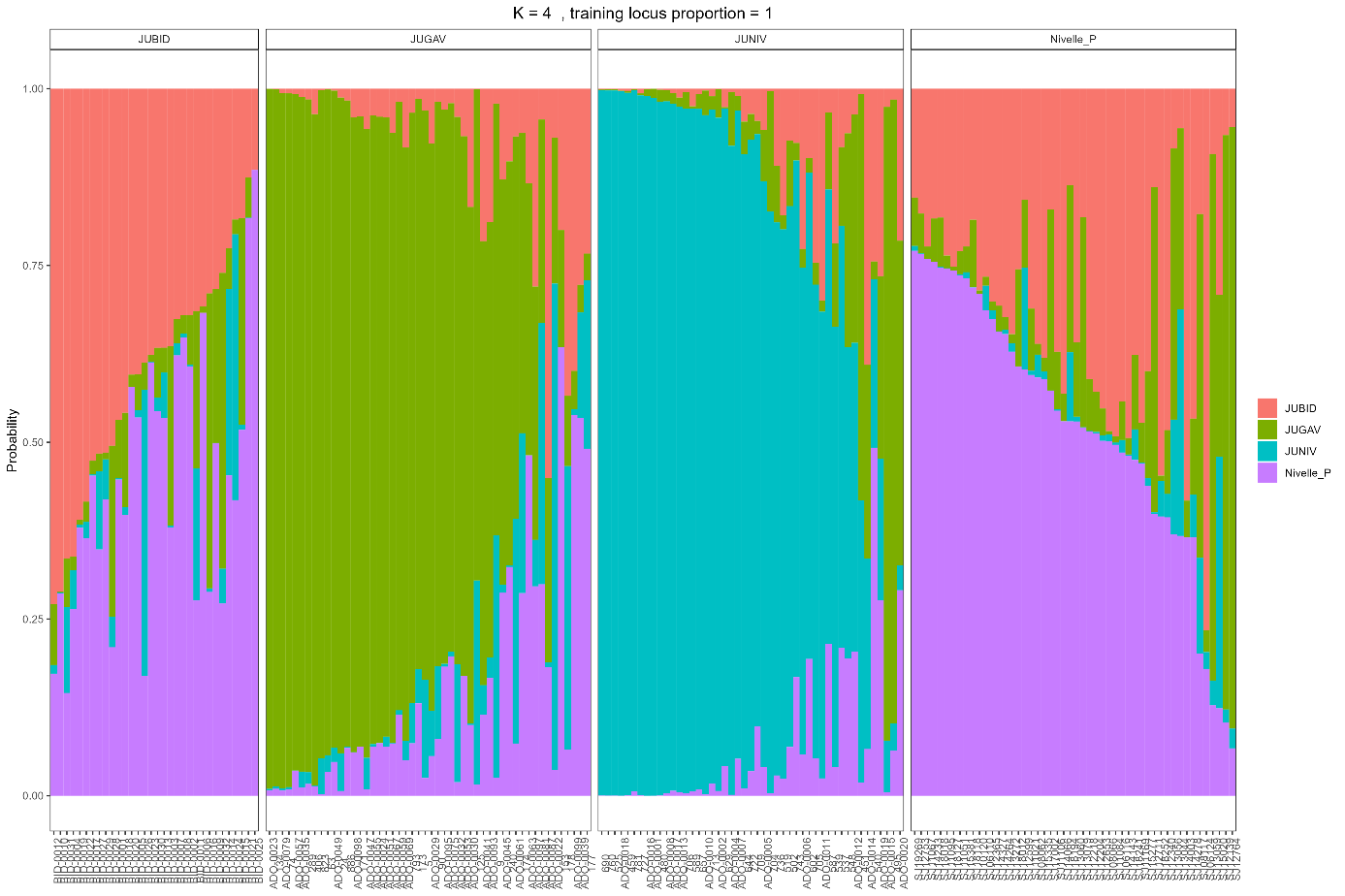

**Fig S2.** **Membership probability of the four source populations used in the study system.** JUBID, JUGAV, JUNIV, and Nivelle_P represent Bidasoa, Gave, Nive and Nivelle populations respectively. Results were estimated via 4-fold cross-validation using overall loci (161 SNPs). Each panel, from left to right, was composed of 32, 50, 47 and 50 individuals respectively.

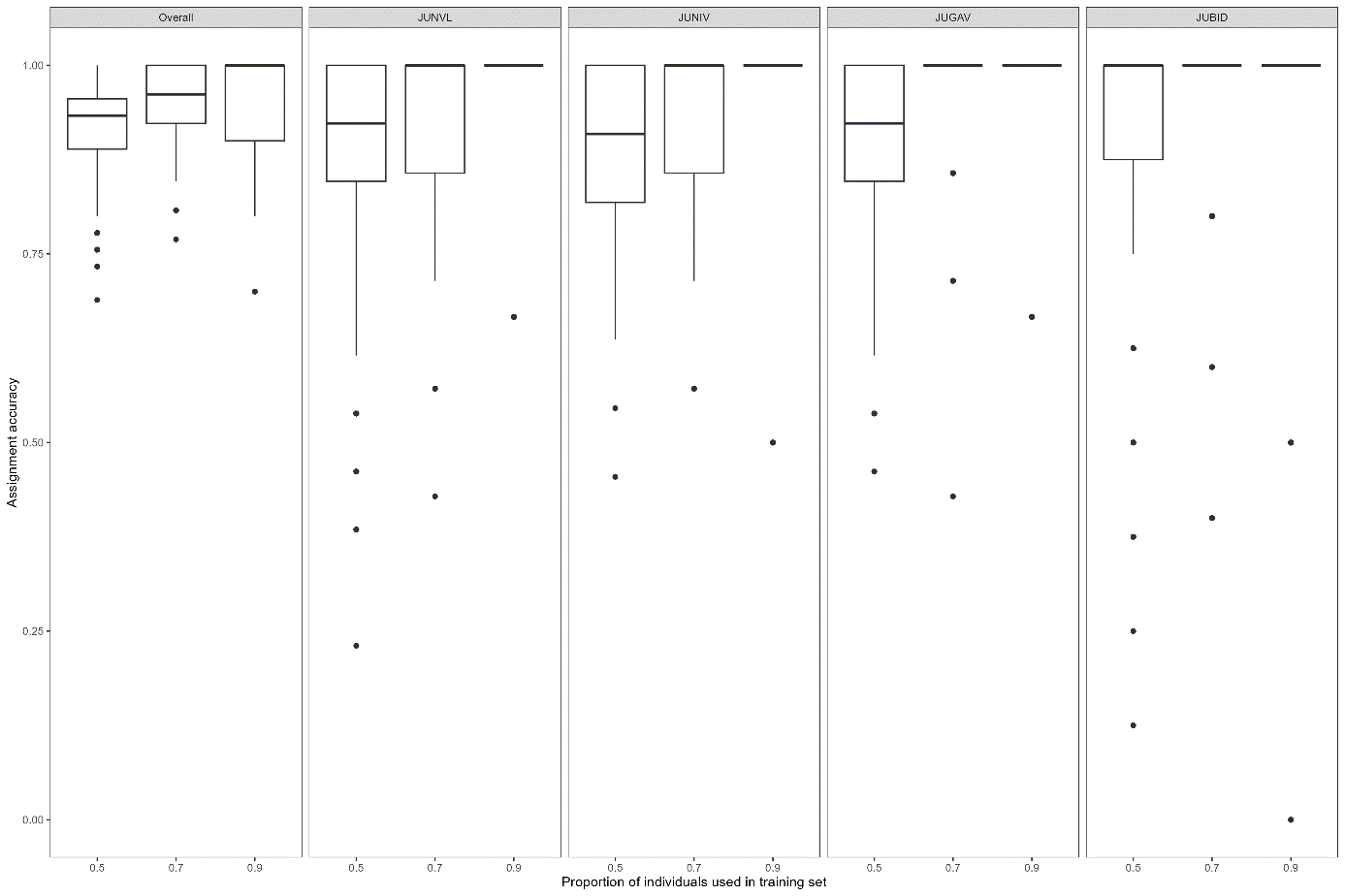

**Fig S3.** **Assignment accuracies estimated after filtering individuals with highest membership probability via Monte-Carlo cross-validation with a linear discriminant analysis (lda) classification method.** We used three levels of training individuals (50%, 70% and 90% of individuals from each population, on x-axis). ‘Overall’ represents the whole dataset, ‘JUNVL’, JUNIV, JUGAV, and JUBID the subset of juveniles from Nivelle, Nive, Gave and Bidasoa populations respectively. We used all available markers for each combination of training individuals.

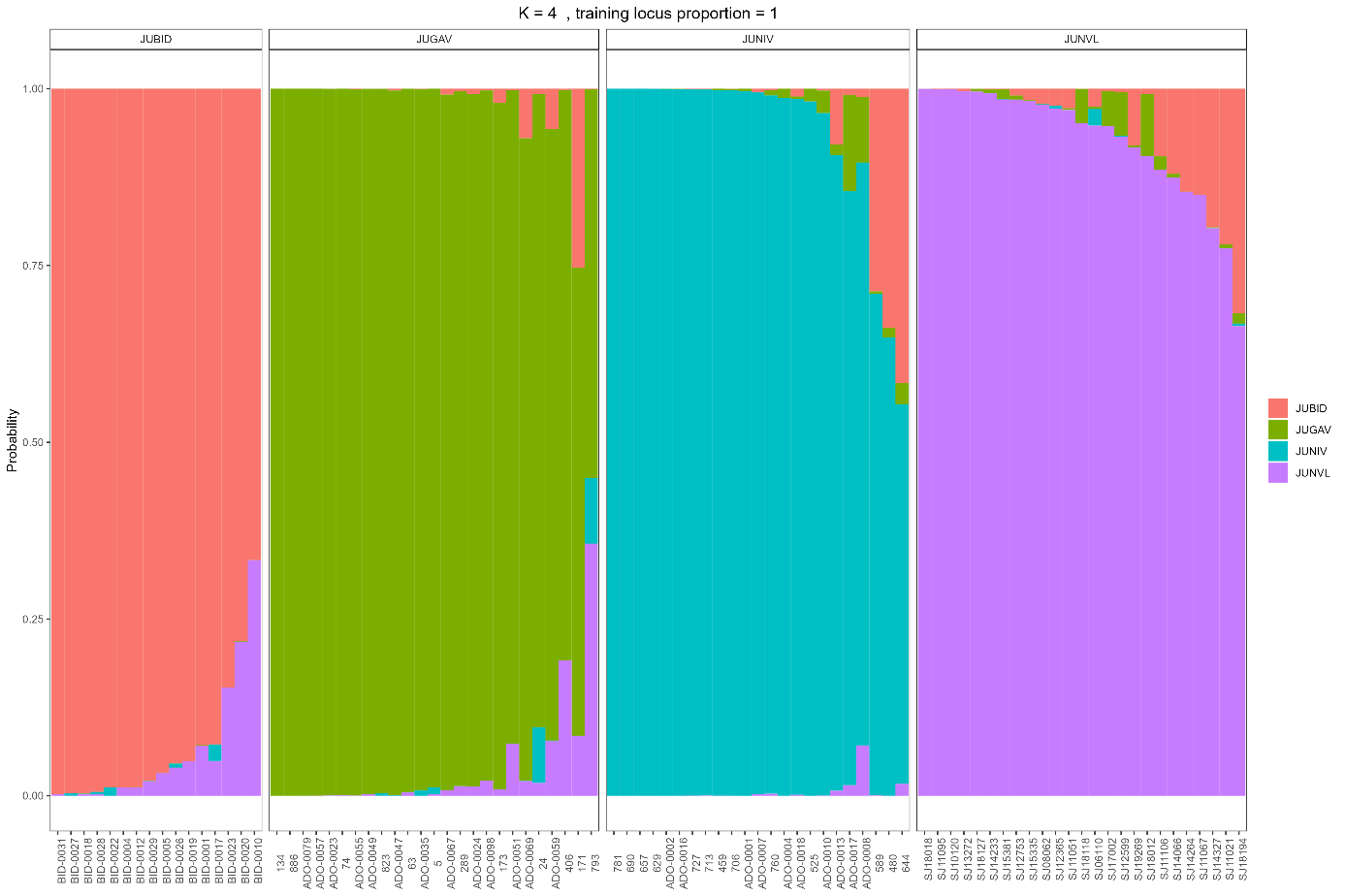

**Fig S4.** **Membership probability estimated after filtering individuals with the highest membership probability of the four populations used in the study system.** JUBID, JUGAV, JUNIV, and JUNVL represent Bidasoa, Gave, Nive and Nivelle populations respectively. Results were estimated via 4-fold cross-validation using overall loci (161 SNPs). Each panel, from left to right, was composed of 25, 23, 16 and 25 individuals respectively.

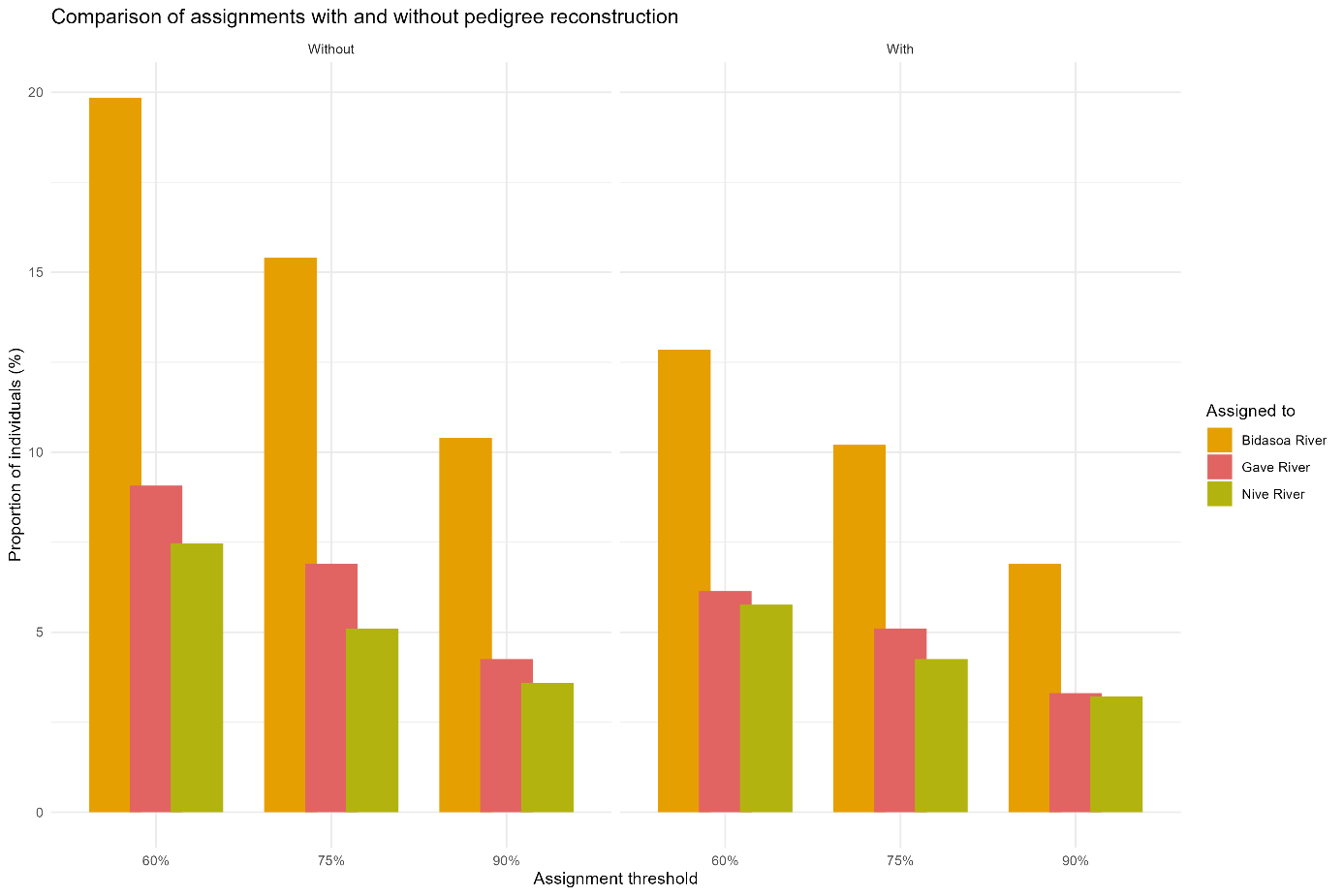

**Fig S5. Comparison of dispersal rates estimated using population assignment tests alone versus population assignment tests combined with pedigree reconstruction.** Dispersal rates were calculated across different assignment probability thresholds (60%, 75%, and 90%) for three river populations: Bidasoa, Gave, and Nive. "Without" refers to dispersal rates estimated using only population assignment tests, while "With" refers to rates estimated after incorporating pedigree information. Higher assignment thresholds result in more conservative estimates but lower sample sizes for rate calculations.

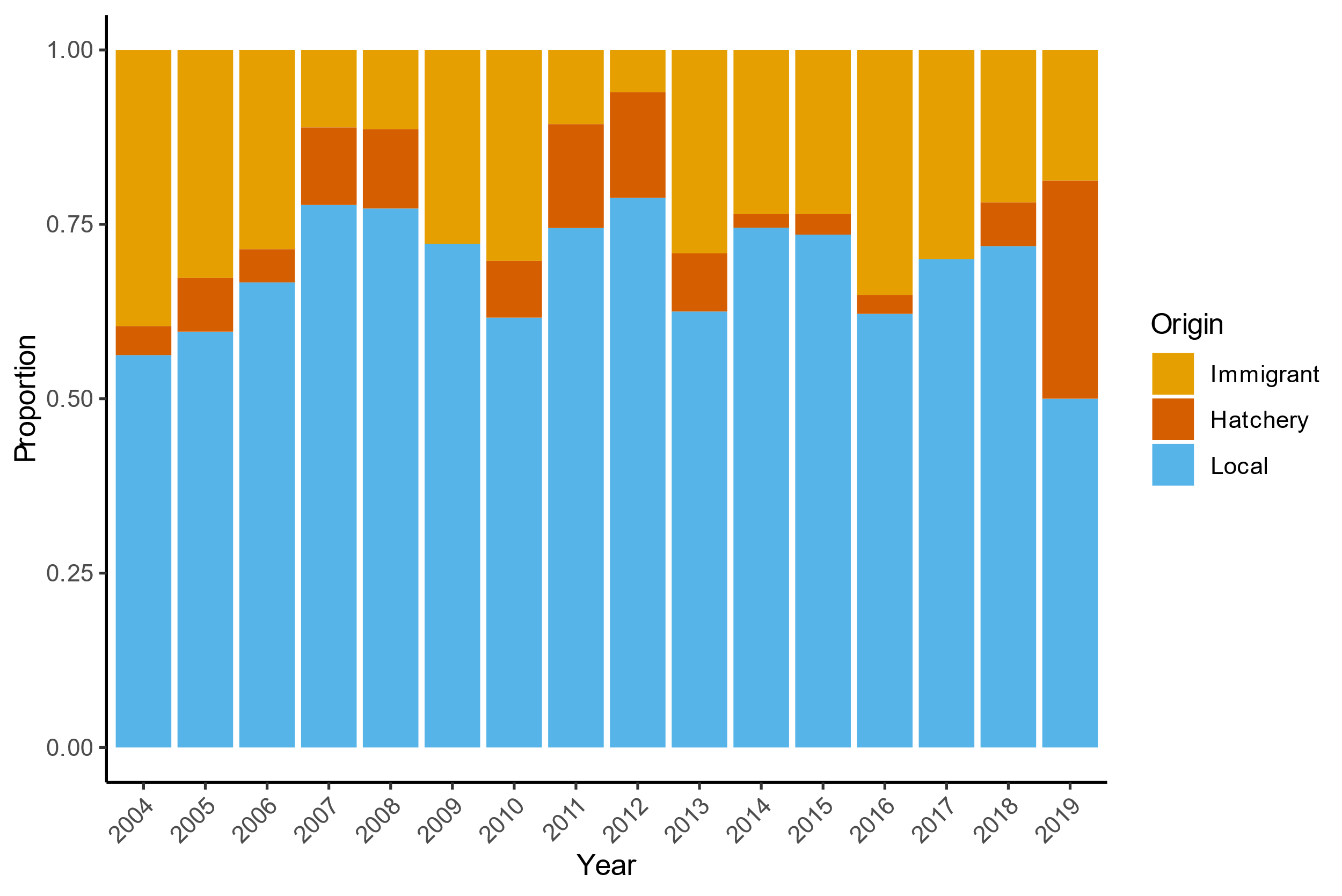

**Fig S6. Temporal variation in the proportion of individuals by origin category from 2004 to 2019.** Each stacked bar represents the relative proportion of individuals classified according to their inferred origin. Category codes: Local refers to individuals assigned to the Nivelle population (residents); Hatchery refers to individuals originating from hatchery populations; Immigrant refers to individuals assigned to wild populations other than their sampling location. Proportions were calculated based on population assignment tests with pedigree correction using individuals with assignment probability q > 75%.

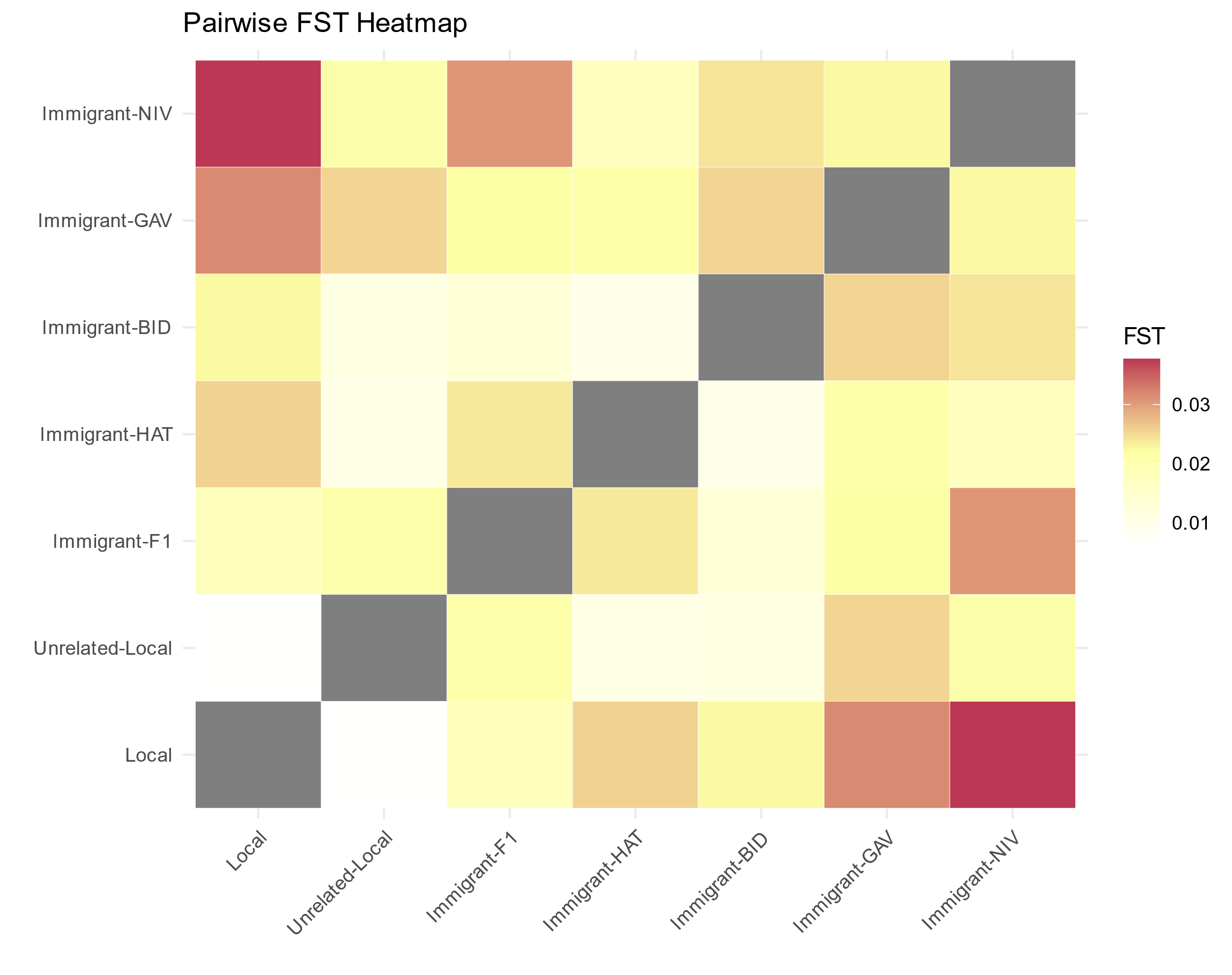

**Fig S7. Pairwise *F_ST_* among populations for 722 adults confidently assigned to their population of origin.** Category codes: Local refers to individuals assigned to the Nivelle population (residents); Local-Unrelated corresponds to individuals assigned to Nivelle source populations but without identified parents in pedigree analysis; Immigrant-F1 represents first-generation immigrants with parents assigned in the Nivelle River but genetically assigned to different source populations; HAT, BID, GAV and NIV represent immigrants from Hatchery, Bidasoa, Gaves and Nive populations, respectively. Heatmap shows FST values with higher values represented by darker shades (scale: 0.01-0.03). All pairwise comparisons were statistically significant (p < 0.05).

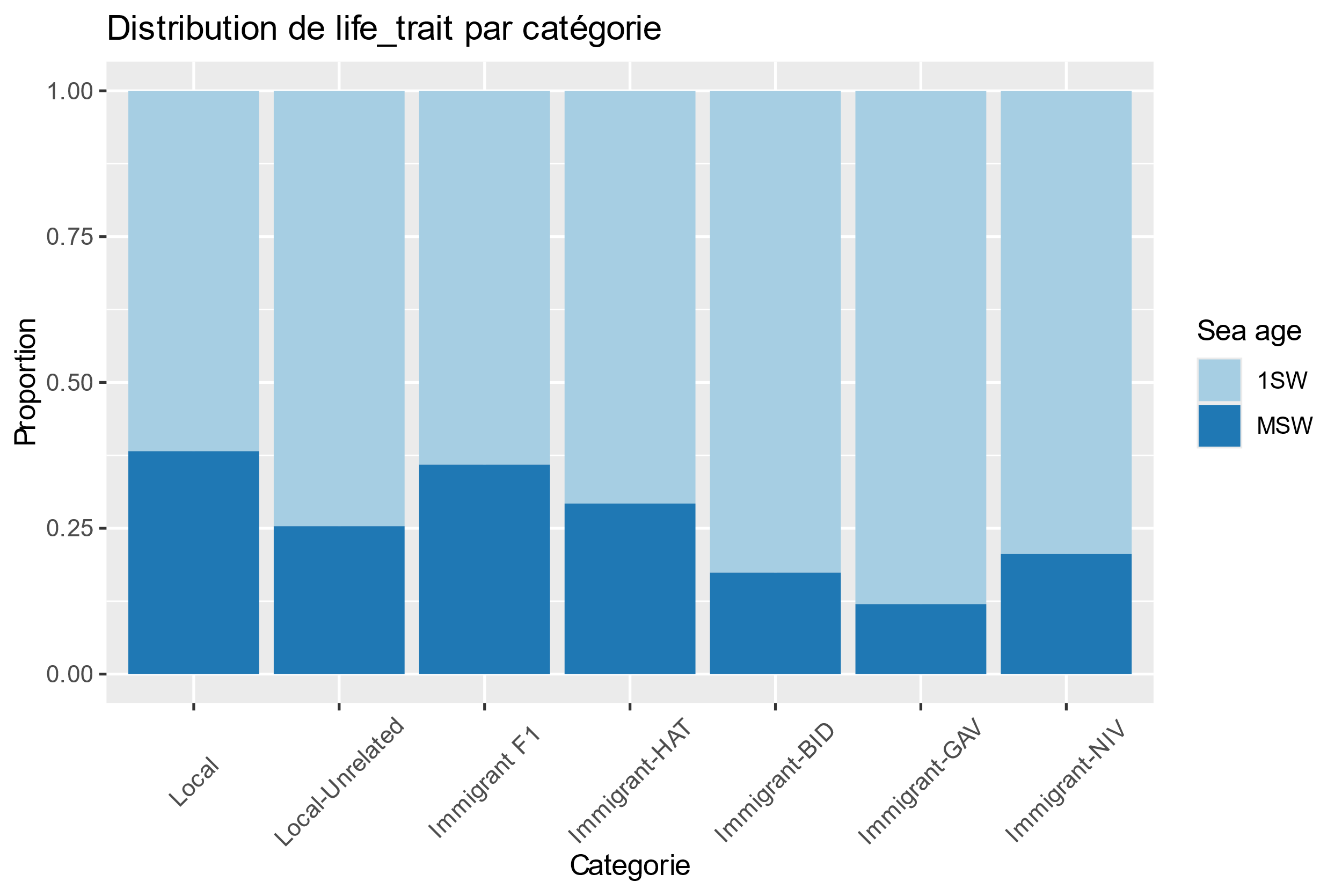
**Fig S8. Sea age proportions (1SW vs MSW) among origin categories for 722 adults confidently assigned to their population of origin.** Category codes: Local refers to individuals assigned to the Nivelle population (residents); Local-Unrelated corresponds to individuals assigned to Nivelle source populations but without identified parents in pedigree analysis; Immigrant-F1 represents first-generation immigrants with parents assigned in the Nivelle River but genetically assigned to different source populations; HAT, BID, GAV and NIV represent immigrants from Hatchery, Bidasoa, Gaves and Nive populations, respectively. Stacked bars show the proportion of 1-sea-winter (1SW, light blue) and multi-sea-winter (MSW, dark blue) individuals within each category. Pairwise comparisons revealed significant differences in 1SW proportions between Local and Immigrant-BID (61.8% vs 82.6%, p = 0.011) and between Local and Immigrant-GAV (61.8% vs 88.0%, p = 0.015) after Bonferroni correction, while all other comparisons were non-significant (p > 0.05).

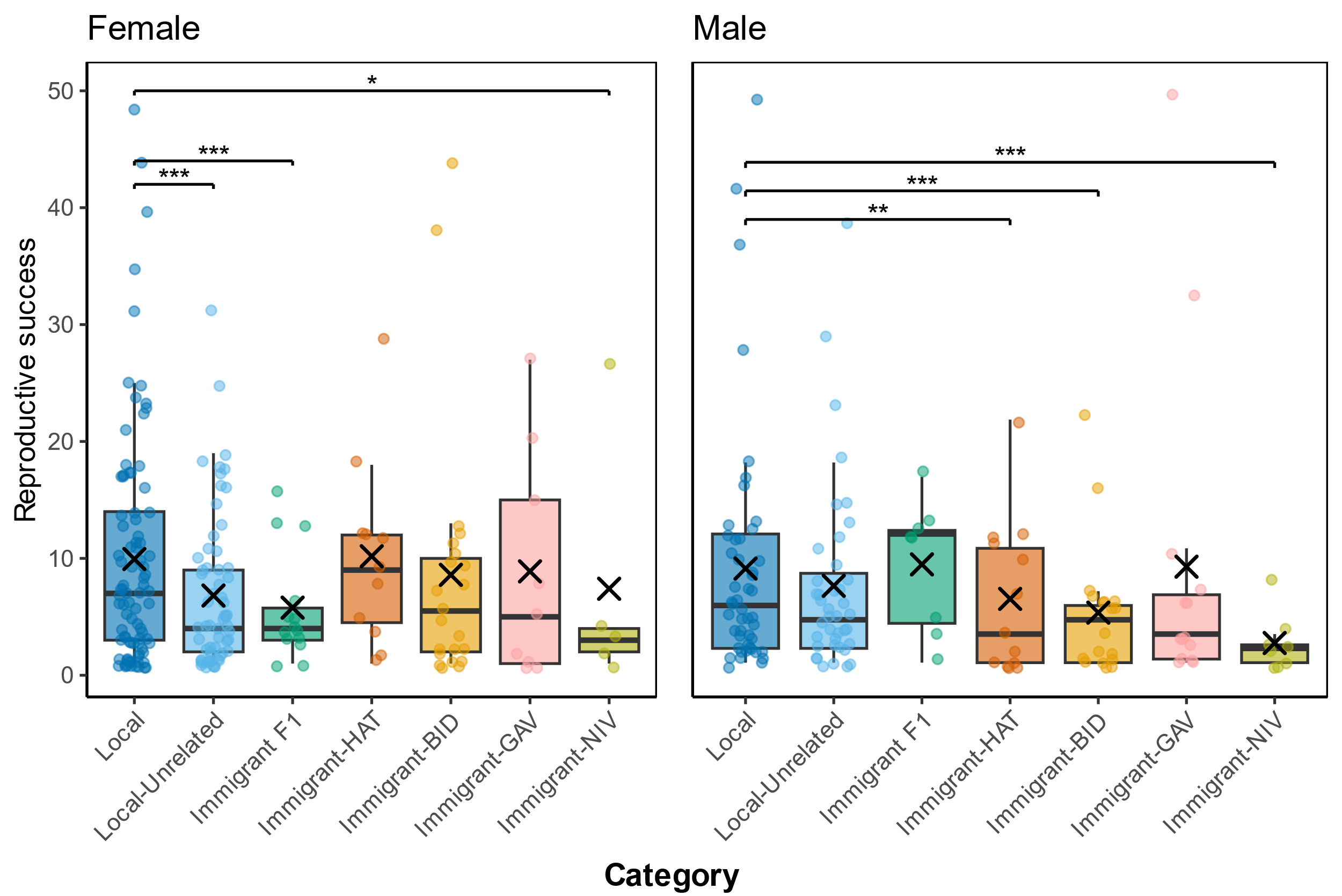

**Fig S9. Reproductive success (number of assigned offspring) among female (left, n = 210) and male (right, n = 149) successful breeders salmon grouped by origin categories.** Large “X” symbols represent group means; small circles show individual data points. Category codes: Local refers to individuals assigned to the Nivelle population (residents); Local-Unrelated corresponds to individuals assigned to Nivelle source populations but without identified parents in pedigree analysis; Immigrant-F1 represents first-generation immigrants with parents assigned in the Nivelle River but genetically assigned to other source populations; HAT, BID, GAV and NIV denote immigrants from Hatchery, Bidasoa, Gaves and Nive populations, respectively. Data points are jittered along the x-axis for clarity. Asterisks indicate statistically significant differences: ***p < 0.001, **p < 0.01, *p < 0.05.

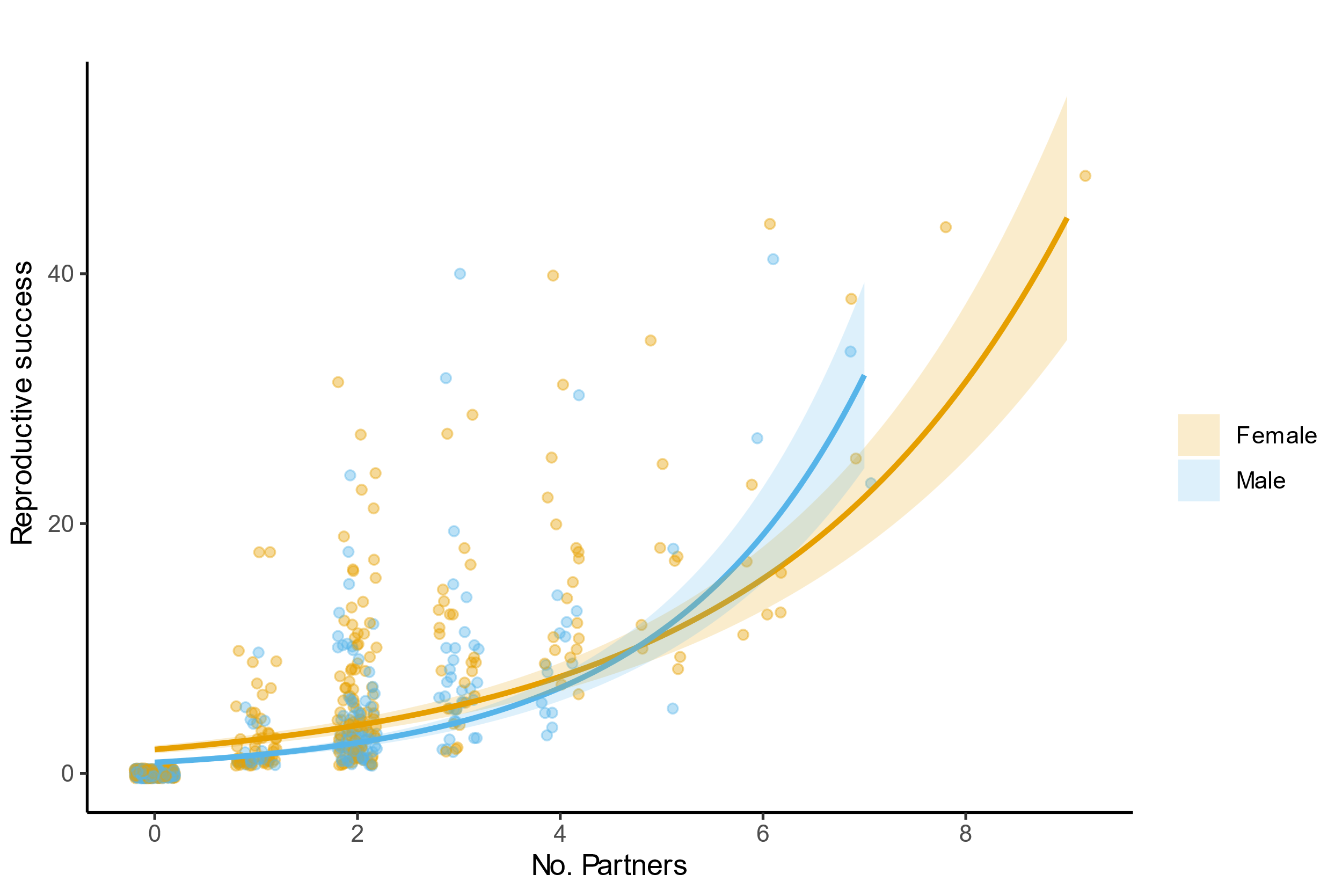

**Fig S10. Bateman gradients showing the relationship between reproductive success and number of partners in 658 adult salmon.** Points represent individual data for female (orange circles) and male (blue circles). Individual data points are jittered along the x-axis for clarity. Lines and shaded areas show fitted relationships (± SE) for each sex.

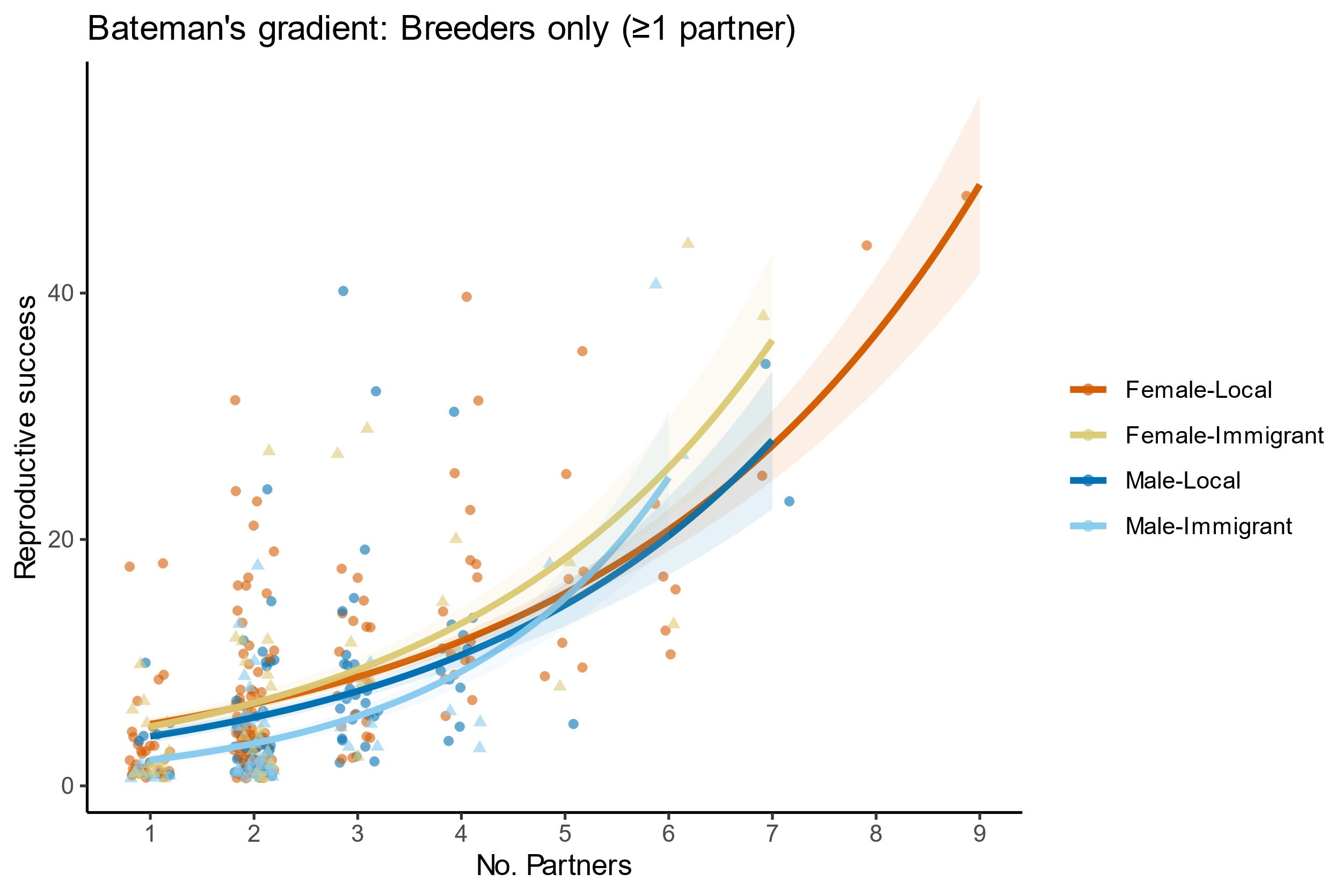

**Fig S11. Bateman gradients showing the relationship between reproductive success and number of partners in 359 successful breeders adult salmon.** Points represent individual data for female locals (orange circles), female immigrants (yellow triangles), male locals (dark blue circles), and male immigrants (light blue triangles). Lines and shaded areas show fitted relationships (± SE) for each sex and origin category.

### Supplemental Tables

**Table S1. Number of genotyped individuals by genotyping method, stage of life, and sampling year.** Individuals were genotyped using microsatellites, SNPs, or both marker types. "Parr" refers to juvenile individuals. N corresponds to the total number of individuals successfully genotyped for each category across all sampling years. Note: For 28 individuals genotyped using SNPs, 7 individuals genotyped using both markers, and 1 individual genotyped with microsatellites, the sampling year was unavailable ("NA" in the dataset).

|  | **microsatellites** | | **SNPs** | | **both** | |
| --- | --- | --- | --- | --- | --- | --- |
|  | **adult** N = 66 | **parr** N = 1,137 | **adult** N = 450 | **parr** N = 1,096 | **adult** N = 648 | **parr** N = 934 |
| year |  |  |  |  |  |  |
| 2003 |  |  | 21 |  |  |  |
| 2004 |  |  | 9 |  | 71 |  |
| 2005 | 1 | 252 | 3 |  | 68 | 34 |
| 2006 |  | 80 | 2 | 4 | 53 | 16 |
| 2007 |  | 98 | 1 |  | 59 | 14 |
| 2008 |  | 201 |  | 1 | 67 | 27 |
| 2009 | 27 | 139 |  | 2 | 24 | 18 |
| 2010 | 7 | 217 | 5 | 4 | 131 | 30 |
| 2011 |  | 149 | 6 | 7 | 65 | 81 |
| 2012 |  | 1 | 5 | 32 | 44 | 252 |
| 2013 | 30 |  | 6 | 10 | 59 | 218 |
| 2014 |  |  | 76 | 28 |  | 244 |
| 2015 |  |  | 54 | 169 |  |  |
| 2016 |  |  | 51 | 187 |  |  |
| 2017 |  |  | 70 | 215 |  |  |
| 2018 |  |  | 42 | 197 |  |  |
| 2019 |  |  | 94 | 217 |  |  |

**Table S2. Summary of parentage assignment runs using FRANz software with different parameter settings.** --Nfmax and --Nmmax correspond to the number of candidate fathers and mothers, respectively. --femrepro and --malerepro specify the age ranges during which females and males can reproduce. "Siblings" indicates whether sibling detection (--fullsibtest) was incorporated in the run. "Genotyping error (%)" refers to the assumed genotyping error rate. "Unrelated (%)" corresponds to individuals where no single genotyped individual could be confidently assigned as a parent. Subsequent analyses were conducted using parameters from Run 9, which excluded sibling testing and yielded the most conservative parentage assignments.

| Run | --femrepro | --malerepro | --Nmmax | --Nfmax | Siblings | Genotyping error (%) | Unrelated (%) | |
| --- | --- | --- | --- | --- | --- | --- | --- | --- |
|  |  |  |  |  |  |  | Parr | Adult |
| 1 | 3:6 | 1:6 | 7600 | 300 | Y | 0 | 32.3 | 61.9 |
| 2 | 3:6 | 3:6 | 7600 | 300 | Y | 0 | 34.6 | 64.1 |
| 3 | 3:6 | 1:6 | 3800 | 300 | Y | 0 | 33.4 | 63.5 |
| 4 | 3:6 | 1:6 | 7600 | 300 | Y | 0.5 | 22.9 | 60.1 |
| 5 | 3:6 | 1:6 | 7600 | 300 | Y | 1 | 22.6 | 54 |
| 6 | 3:6 | 1:6 | 7600 | 300 | Y | 3 | 24.7 | 53.5 |
| 7 | 3:6 | 1:6 | 7600 | 300 | Y | 6 | 32.7 | 58.5 |
| 8 | 3:6 | 3:6 | 7600 | 300 | Y | 1 | 24.5 | 61.5 |
| **9** | 3:6 | 1:6 | 7600 | 300 | N | 1 | 15.4 | 46.9 |

**Table S3. Population assignment results using Naïve Bayes classification corrected with pedigree information.** Population assignment tests were performed on adult individuals, with each individual assigned to a predicted source population based on membership probability (q). Two probability thresholds were applied: q > 50% and q > 75%. Assignment success was evaluated using population-only genetic information and after incorporating pedigree correction. "Gave Juveniles," "Nive Juveniles," "Bidasoa Juveniles," and "Nivelle Juveniles" refer to the predicted natal populations of assigned adults. Numbers in parentheses represent the percentage of total assignments for each category. Final dispersal rates were computed using an integrative approach based on the total adult sample size (N = 1058) and are highlighted in bold.

|  | ***q* > 50%** | | ***q* > 75%** | |
| --- | --- | --- | --- | --- |
|  | **Total: 1058 individuals** | | **Total: 1058 individuals** | |
|  | **Only Population assignment tests** | **Pedigree correction** | **Only Population assignment tests** | **Pedigree correction** |
| **Assigned to** |  |  |  |  |
| Gave Juveniles | 114 (10.8%) | 77 (7.3%) | 73 (6,9%) | 54 (**5.1%)** |
| Nive Juveniles | 93 (8.8%) | 70 (6.62%) | 54 (5.1%) | 45 (**4.2%)** |
| Bidasoa Juveniles | 256 (24.2%) | 166 (15.7%) | 163 (15.4%) | 108 (**10.2%)** |
| Nivelle Juveniles | 595 (56.2%) | 311 (29.4%) | 478 (45.2%) | 253 (24%) |
| **Subset of individuals** | 1058 | 624 | 768 | 460 |

**Table S4. Summary of generalized linear models (GLMs) testing phenotypic differences between immigrant and local Atlantic salmon adults** (n = 722). We used GLMs with binomial error distribution and logit link function with a binary response variable (local = 0, immigrant = 1). Models tested effects of sea age at maturity (1SW vs MSW), time spent in river (river age), and sex on dispersal status. To avoid multicollinearity, separate models were fitted for each predictor variable. Sex effects were analysed separately for one-sea-winter (1SW) and multi-sea-winter (MSW) individuals. An offset term based on the logarithm of annual sampling effort was included to control for yearly variation in sampling intensity. N = sample size, 95% CI = 95% Confidence Interval, P = p-value. Reference categories: sea age (1SW), river age (1 year), sex (Female).

Table S4.1 – Sea age and dispersal

| **Term** | **N** | **Parameter estimate** | **95% CI** | **P** |
| --- | --- | --- | --- | --- |
| **(Intercept)** | 722 | -4.4 | -4.6, -4.3 | <0.001 |
| **Sea age** |  |  |  |  |
| 1SW | 519 | — | — |  |
| MSW | 203 | -0.53 | -0.91, -0.16 | 0.006 |

Table S4.2 – River age and dispersal

| **Term** | **N** | **Parameter estimate** | **95% CI** | **P** |
| --- | --- | --- | --- | --- |
| **(Intercept)** | 717 | -4.6 | -4.8, -4.4 | <0.001 |
| **Freshwater age** |  |  |  |  |
| 1 | 597 | — | — |  |
| 2 | 120 | 0.22 | -0.20, 0.62 | 0.3 |

Table S4.3 – Sex and dispersal

| **Term** | **N** | **Parameter estimate** | **95% CI** | **P** |
| --- | --- | --- | --- | --- |
| **(Intercept)** | 722 | -4.8 | -5.0, -4.6 | <0.001 |
| **Sex** |  |  |  |  |
| F | 377 | — | — |  |
| M | 345 | 0.45 | 0.14, 0.77 | 0.005 |

Table S4.4 – Only 1SW: Sex and dispersal

| **Term** | **N** | **Parameter estimate** | **95% CI** | **P** |
| --- | --- | --- | --- | --- |
| **(Intercept)** | 519 | -4.6 | -4.9, -4.3 | <0.001 |
| **Sex** |  |  |  |  |
| F | 203 | — | — |  |
| M | 316 | 0.19 | -0.18, 0.57 | 0.3 |

Table S4.5 – Only MSW: Sex and dispersal

| **Term** | **N** | **Parameter estimate** | **95% CI** | **P** |
| --- | --- | --- | --- | --- |
| **(Intercept)** | 203 | -5.1 | -5.5, -4.8 | <0.001 |
| **Sex** |  |  |  |  |
| F | 174 | — | — |  |
| M | 29 | 0.98 | 0.13, 1.8 | 0.021 |

**Table S5. Summary of generalized linear models (GLMs) testing effects of phenotypic traits on reproductive success (RS) in Atlantic salmon adults (n = 658).** Zero-inflated models tested effects of body size (log-transformed mass in grams), sea age at maturity (1SW vs MSW), and sex on reproductive success. To avoid multicollinearity, separate models were fitted for each predictor variable as a fixed effect. Body size effects were analysed both globally and specifically for one-sea-winter (1SW) individuals (n = 463). An offset term based on the logarithm of offspring sampled in each parent's sampling year was included in both model components to control for annual variation in sampling effort. N = sample size, IRR = Incidence Rate Ratio, 95% CI = 95% Confidence Interval, P = p-value. Reference categories: sea age (1SW), sex (Female).

Table S5.1 – Mass and RS

| **Term** | **Parameter estimate** | **95% CI***^1^* | **P** |
| --- | --- | --- | --- |
| (Intercept) | -8.9 | -9.6, -8.1 | <0.001 |
| log(mass_g) | 0.69 | 0.60, 0.78 | <0.001 |
| Zero-inflation | -5.5 | -5.7, -5.4 | <0.001 |

Table S5.2 – Only 1SW: Mass and RS

| **Term** | **Parameter estimate** | **95% CI***^1^* | **P** |
| --- | --- | --- | --- |
| (Intercept) | -9.2 | -11, -7.9 | <0.001 |
| log(mass_g) | 0.74 | 0.56, 0.91 | <0.001 |
| Zero-inflation | -5.4 | -5.6, -5.2 | <0.001 |
| *^1^*CI = Confidence Interval | | | |

Table S5.3 – Sea age and RS

| **Term** | **Parameter estimate** | **95% CI***^1^* | **P** |
| --- | --- | --- | --- |
| (Intercept) | -3.5 | -3.6, -3.5 | <0.001 |
| Sea age |  |  |  |
| 1SW | — | — |  |
| MSW | 0.48 | 0.41, 0.56 | <0.001 |
| Zero-inflation | -5.5 | -5.7, -5.4 | <0.001 |
| *^1^*CI = Confidence Interval | | | |

Table S5.4 – Sex and RS

| **Term** | **Parameter estimate** | **95% CI***^1^* | **P** |
| --- | --- | --- | --- |
| (Intercept) | -3.2 | -3.3, -3.2 | <0.001 |
| Sex |  |  |  |
| F | — | — |  |
| M | -0.32 | -0.40, -0.24 | <0.001 |
| Zero-inflation | -5.5 | -5.7, -5.4 | <0.001 |
| *^1^*CI = Confidence Interval | | | |

**Table S6. Summary of generalized linear models (GLMs) testing effects of origin on reproductive success (RS) in Atlantic salmon adults (n = 658).** Zero-inflated models tested effects of dispersal status (local vs immigrant) on RS analysed globally and separately by sex (Female, n = 361; Male, n = 297) and sea age at maturity (1SW, n = 363; MSW, n = 195). An offset term based on the logarithm of offspring sampled in each parent's sampling year was included in both model components to control for annual variation in sampling effort. N = sample size, IRR = Incidence Rate Ratio, 95% CI = 95% Confidence Interval, P = p-value. Reference category: local (0).

Table S6.1 – Origin and RS

| **Term** | **Parameter estimate** | **95% CI***^1^* | **P** |
| --- | --- | --- | --- |
| (Intercept) | -3.3 | -3.3, -3.2 | <0.001 |
| Origin |  |  |  |
| 0 | — | — |  |
| 1 | -0.16 | -0.25, -0.07 | <0.001 |
| Zero-inflation | -5.5 | -5.7, -5.4 | <0.001 |
| *^1^*CI = Confidence Interval | | | |

Table S6.2 – Only females: origin and RS

| **Term** | **Parameter estimate** | **95% CI***^1^* | **P** |
| --- | --- | --- | --- |
| (Intercept) | -3.2 | -3.3, -3.2 | <0.001 |
| Origin |  |  |  |
| 0 | — | — |  |
| 1 | 0.05 | -0.06, 0.16 | 0.4 |
| Zero-inflation | -5.6 | -5.8, -5.4 | <0.001 |
| *^1^*CI = Confidence Interval | | | |

Table S6.3 – Only males: origin and RS

| **Term** | **Parameter estimate** | **95% CI***^1^* | **P** |
| --- | --- | --- | --- |
| (Intercept) | -3.4 | -3.5, -3.3 | <0.001 |
| Origin |  |  |  |
| 0 | — | — |  |
| 1 | -0.35 | -0.50, -0.21 | <0.001 |
| Zero-inflation | -5.4 | -5.6, -5.1 | <0.001 |
| *^1^*CI = Confidence Interval | | | |

Table S6.4 – Only 1SW: origin and RS

| **Term** | **Parameter estimate** | **95% CI***^1^* | **P** |
| --- | --- | --- | --- |
| (Intercept) | -3.5 | -3.5, -3.4 | <0.001 |
| Origin |  |  |  |
| 0 | — | — |  |
| 1 | -0.19 | -0.31, -0.08 | <0.001 |
| Zero-inflation | -5.4 | -5.5, -5.2 | <0.001 |
| *^1^*CI = Confidence Interval | | | |

Table S6.5 – Only MSW: origin and RS

| **Term** | **Parameter estimate** | **95% CI***^1^* | **P** |
| --- | --- | --- | --- |
| (Intercept) | -3.1 | -3.1, -3.0 | <0.001 |
| Origin |  |  |  |
| 0 | — | — |  |
| 1 | 0.20 | 0.05, 0.34 | 0.007 |
| Zero-inflation | -5.9 | -6.2, -5.6 | <0.001 |
| *^1^*CI = Confidence Interval | | | |

**Table S7. Summary of generalized linear models (GLMs) testing effects of population origin categories on reproductive success in Atlantic salmon adults (n = 658).** Zero-inflated models tested effects of population origin categories (Local_pur = Local, Local_without_parents = Local_Unrelated, Descendant_immigrant = Immigrant-F1, Hatchery_disperser = Immigrant-HAT, Immigrant-BID, Immigrant-GAV, Immigrant-NIV) on reproductive success analysed globally, separately by sex (Female, n = 361; Male, n = 297), for one-sea-winter individuals only, for multi-sea-winter individuals only, and stratified by sea age class. An offset term based on the logarithm of offspring sampled in each parent’s sampling year was included in both model components to control for annual variation in sampling effort. N = sample size, IRR = Incidence Rate Ratio, 95% CI = 95% Confidence Interval, P = p-value. Reference category: Local_pur (Local).

Table S7.1 – Origin categories, sex and RS

| **Term** | **Parameter estimate** | **95% CI***^1^* | **P** |
| --- | --- | --- | --- |
| (Intercept) | -3.1 | -3.2, -3.0 | <0.001 |
| Origin category |  |  |  |
| local_pur | — | — |  |
| local_without_parents | -0.20 | -0.32, -0.09 | <0.001 |
| descendant_immigrant | -0.59 | -0.82, -0.36 | <0.001 |
| hatchery_disperser | 0.14 | -0.05, 0.34 | 0.2 |
| pure_bid_disperser | -0.11 | -0.26, 0.05 | 0.2 |
| pure_gav_disperser | -0.03 | -0.26, 0.20 | 0.8 |
| pure_niv_disperser | -0.40 | -0.74, -0.06 | 0.021 |
| Sex |  |  |  |
| F | — | — |  |
| M | -0.28 | -0.40, -0.16 | <0.001 |
| Origin category * Sex |  |  |  |
| local_without_parents * M | 0.12 | -0.08, 0.33 | 0.2 |
| descendant_immigrant * M | 0.69 | 0.34, 1.0 | <0.001 |
| hatchery_disperser * M | -0.50 | -0.82, -0.18 | 0.002 |
| pure_bid_disperser * M | -0.41 | -0.71, -0.11 | 0.007 |
| pure_gav_disperser * M | 0.03 | -0.29, 0.34 | 0.9 |
| pure_niv_disperser * M | -0.87 | -1.5, -0.26 | 0.006 |
| Zero-inflation | -5.5 | -5.7, -5.4 | <0.001 |
| *^1^*CI = Confidence Interval Table S7.2 – Separated by sex: Origin categories and RS \| **RS / FEMALES ** \| \| \| **RS / MALES ** \| \|; **Term** \| **Parameter estimate** \| **95% CI***^1^* \| **P** \| **Parameter estimate** \| **95% CI***^1^* \| **P**; (Intercept) \| -3.1 \| -3.2, -3.0 \| <0.001 \| -3.4 \| -3.5, -3.3 \| <0.001; Origin category \| \| \| \| \| \|; local_pur \| — \| — \| \| — \| — \|; local_without_parents \| -0.21 \| -0.32, -0.09 \| <0.001 \| -0.08 \| -0.24, 0.08 \| 0.3; descendant_immigrant \| -0.59 \| -0.82, -0.36 \| <0.001 \| 0.10 \| -0.17, 0.37 \| 0.5; hatchery_disperser \| 0.14 \| -0.05, 0.34 \| 0.2 \| -0.36 \| -0.61, -0.10 \| 0.007; pure_bid_disperser \| -0.11 \| -0.27, 0.05 \| 0.2 \| -0.51 \| -0.76, -0.26 \| <0.001; pure_gav_disperser \| -0.03 \| -0.26, 0.20 \| 0.8 \| 0.00 \| -0.22, 0.22 \| >0.9; pure_niv_disperser \| -0.41 \| -0.75, -0.06 \| 0.021 \| -1.3 \| -1.8, -0.75 \| <0.001; Zero-inflation \| -5.6 \| -5.9, -5.4 \| <0.001 \| -5.4 \| -5.6, -5.1 \| <0.001; *^1^*CI = Confidence Interval \| \| \| \| \| \| Table S7.3 – Separated by sex / Only 1SW: Origin categories and RS \| **RS / FEMALES / 1SW ** \| \| \| **RS / MALES / 1SW ** \| \|; **Term** \| **Parameter estimate** \| **95% CI***^1^* \| **P** \| **Parameter estimate** \| **95% CI***^1^* \| **P**; (Intercept) \| -3.4 \| -3.5, -3.3 \| <0.001 \| -3.3 \| -3.5, -3.2 \| <0.001; Origin category \| \| \| \| \| \|; local_pur \| — \| — \| \| — \| — \|; local_without_parents \| -0.28 \| -0.49, -0.08 \| 0.007 \| -0.12 \| -0.29, 0.05 \| 0.2; descendant_immigrant \| -0.54 \| -0.93, -0.16 \| 0.006 \| 0.22 \| -0.06, 0.51 \| 0.12; hatchery_disperser \| 0.53 \| 0.28, 0.79 \| <0.001 \| -0.60 \| -0.90, -0.30 \| <0.001; pure_bid_disperser \| -0.49 \| -0.76, -0.22 \| <0.001 \| -0.49 \| -0.74, -0.23 \| <0.001; pure_gav_disperser \| 0.20 \| -0.08, 0.48 \| 0.2 \| -0.04 \| -0.27, 0.19 \| 0.7; pure_niv_disperser \| -1.8 \| -2.8, -0.80 \| <0.001 \| -1.7 \| -2.4, -1.1 \| <0.001; Zero-inflation \| -5.6 \| -5.9, -5.3 \| <0.001 \| -5.3 \| -5.5, -5.0 \| <0.001; *^1^*CI = Confidence Interval \| \| \| \| \| \| Table S7.4 – Separated by sex / Only MSW: Origin categories and RS \| **RS / FEMALES / MSW ** \| \| \| **RS / MALES / MSW ** \| \|; **Term** \| **Parameter estimate** \| **95% CI***^1^* \| **P** \| **Parameter estimate** \| **95% CI***^1^* \| **P**; (Intercept) \| -2.9 \| -3.0, -2.9 \| <0.001 \| -3.6 \| -3.8, -3.3 \| <0.001; Origin category \| \| \| \| \| \|; local_pur \| — \| — \| \| — \| — \|; local_without_parents \| -0.06 \| -0.20, 0.09 \| 0.5 \| 0.06 \| -0.52, 0.64 \| 0.8; descendant_immigrant \| -0.55 \| -0.84, -0.27 \| <0.001 \| -0.80 \| -1.8, 0.20 \| 0.12; hatchery_disperser \| -0.18 \| -0.53, 0.17 \| 0.3 \| 0.82 \| 0.29, 1.3 \| 0.002; pure_bid_disperser \| 0.57 \| 0.37, 0.76 \| <0.001 \| -1.3 \| -2.4, -0.12 \| 0.030; pure_gav_disperser \| 0.28 \| -0.24, 0.79 \| 0.3 \| 0.10 \| -0.63, 0.83 \| 0.8; pure_niv_disperser \| -0.04 \| -0.39, 0.32 \| 0.8 \| -0.22 \| -0.95, 0.52 \| 0.6; Zero-inflation \| -5.8 \| -6.1, -5.5 \| <0.001 \| -6.6 \| -7.6, -5.7 \| <0.001; *^1^*CI = Confidence Interval \| \| \| \| \| \| Table S7.5 – Separated by sea age class: individual categories and RS \| **RS / 1SW ** \| \| \| **RS / MSW ** \| \|; **Term** \| **Parameter estimate** \| **95% CI***^1^* \| **P** \| **Parameter estimate** \| **95% CI***^1^* \| **P**; (Intercept) \| -3.4 \| -3.5, -3.3 \| <0.001 \| -3.0 \| -3.1, -3.0 \| <0.001; Origin category \| \| \| \| \| \|; local_pur \| — \| — \| \| — \| — \|; local_without_parents \| -0.18 \| -0.32, -0.05 \| 0.006 \| -0.01 \| -0.15, 0.13 \| >0.9; descendant_immigrant \| -0.11 \| -0.34, 0.11 \| 0.3 \| -0.58 \| -0.86, -0.31 \| <0.001; hatchery_disperser \| -0.06 \| -0.26, 0.13 \| 0.5 \| 0.02 \| -0.27, 0.30 \| >0.9; pure_bid_disperser \| -0.49 \| -0.68, -0.30 \| <0.001 \| 0.41 \| 0.22, 0.60 \| <0.001; pure_gav_disperser \| 0.06 \| -0.12, 0.24 \| 0.5 \| -0.01 \| -0.42, 0.39 \| >0.9; pure_niv_disperser \| -1.8 \| -2.3, -1.2 \| <0.001 \| -0.18 \| -0.50, 0.14 \| 0.3; Zero-inflation \| -5.4 \| -5.6, -5.2 \| <0.001 \| -5.9 \| -6.2, -5.6 \| <0.001; *^1^*CI = Confidence Interval \| \| \| \| \| \| | | | |

**Table S8. Summary of generalized linear models (GLMs) testing effects of origin, sea age, sex, and origin categories on mating success in Atlantic salmon adults (n = 658).** Zero-inflated models tested effects of origin (0 = local, 1 = immigrant), sea age and sex separately on mating success; effects of origin categories (Local_pur = Local, Local_without_parents = Local_Unrelated, Descendant_immigrant = Immigrant-F1, Hatchery_disperser = Immigrant-HAT, Immigrant-BID, Immigrant-GAV, Immigrant-NIV) on mating success, separated by sex (Female, n = 361; Male, n = 297) and then by sea age (1SW, n = 463; MSW, n = 195). An offset term based on the logarithm of offspring sampled in each parent’s sampling year was included in both model components to control for annual variation in sampling effort. N = sample size, IRR = Incidence Rate Ratio, 95% CI = 95% Confidence Interval, P = p-value. Reference category: Local_pur (Local).

Table S8.1 – Origin and mating success

| **Term** | **N** | **Parameter estimate** | **95% CI***^1^* | **P** |
| --- | --- | --- | --- | --- |
| **(Intercept)** | 658 | -3.0 | -3.1, -2.9 | <0.001 |
| **Origin** |  |  |  |  |
| 0 | 449 | — | — |  |
| 1 | 209 | -0.20 | -0.38, -0.03 | 0.020 |
| **Zero-inflation** | 658 | -0.48 | -0.67, -0.30 | <0.001 |
| *^1^*CI = Confidence Interval | | | | |

Table S8.2 – Sea age and mating success

| **Term** | **N** | **Parameter estimate** | **95% CI***^1^* | **P** |
| --- | --- | --- | --- | --- |
| **(Intercept)** | 658 | -3.2 | -3.3, -3.1 | <0.001 |
| **Sea age** |  |  |  |  |
| 1SW | 463 | — | — |  |
| MSW | 195 | 0.42 | 0.27, 0.57 | <0.001 |
| **Zero-inflation** | 658 | -0.52 | -0.71, -0.32 | <0.001 |
| *^1^*CI = Confidence Interval | | | | |

Table S8.3 – Sex and mating success

| **Term** | **N** | **Parameter estimate** | **95% CI***^1^* | **P** |
| --- | --- | --- | --- | --- |
| **(Intercept)** | 658 | -3.0 | -3.1, -2.9 | <0.001 |
| **sex** |  |  |  |  |
| F | 361 | — | — |  |
| M | 297 | -0.08 | -0.24, 0.07 | 0.3 |
| **Zero-inflation** | 658 | -0.47 | -0.66, -0.29 | <0.001 |
| *^1^*CI = Confidence Interval | | | | |

Table S8.4 – Separated by sex: origin categories and mating success

|  | **Mating success Females** | | | | **Mating success Males** | | | |
| --- | --- | --- | --- | --- | --- | --- | --- | --- |
| **Term** | **N** | **Parameter estimate** | **95% CI***^1^* | **P** | **N** | **Parameter estimate** | **95% CI***^1^* | **P** |
| **(Intercept)** | 361 | -2.8 | -3.0, -2.7 | <0.001 | 297 | -2.9 | -3.1, -2.7 | <0.001 |
| **Categorie** |  |  |  |  |  |  |  |  |
| local_pur | 132 | — | — |  | 78 | — | — |  |
| local_without_parents | 108 | -0.42 | -0.66, -0.18 | <0.001 | 92 | -0.28 | -0.59, 0.02 | 0.066 |
| descendant_immigrant | 22 | -0.43 | -0.83, -0.03 | 0.037 | 17 | 0.09 | -0.43, 0.61 | 0.7 |
| hatchery_disperser | 23 | -0.10 | -0.55, 0.34 | 0.6 | 22 | -0.45 | -0.89, 0.00 | 0.051 |
| pure_bid_disperser | 44 | -0.32 | -0.66, 0.01 | 0.060 | 42 | -0.63 | -1.1, -0.20 | 0.004 |
| pure_gav_disperser | 20 | -0.56 | -1.1, -0.01 | 0.046 | 28 | 0.00 | -0.38, 0.38 | >0.9 |
| pure_niv_disperser | 12 | -0.85 | -1.7, 0.01 | 0.053 | 18 | -0.22 | -0.79, 0.36 | 0.5 |
| **Zero-inflation** | 361 | -0.69 | -0.97, 0.42 | <0.001 | 297 | -0.32 | -0.60, -0.04 | 0.024 |
| *^1^*CI = Confidence Interval | | | | | | | | |

Table S8.5 – Separated by sea age classes: Individual categories and mating success

|  | **Mating success 1SW** | | | | **Mating success MSW** | | | |
| --- | --- | --- | --- | --- | --- | --- | --- | --- |
| **Term** | **N** | **Parameter estimate** | **95% CI***^1^* | **P** | **N** | **Parameter estimate** | **95% CI***^1^* | **P** |
| **(Intercept)** | 463 | -3.2 | -3.4, -3.1 | <0.001 | 195 | -2.9 | -3.1, -2.8 | <0.001 |
| **Categorie** |  |  |  |  |  |  |  |  |
| local_pur | 125 | — | — |  | 85 | — | — |  |
| local_without_parents | 148 | -0.31 | -0.49, -0.12 | 0.001 | 52 | -0.18 | -0.41, 0.06 | 0.14 |
| descendant_immigrant | 25 | -0.23 | -0.55, 0.10 | 0.2 | 14 | -0.17 | -0.57, 0.24 | 0.4 |
| hatchery_disperser | 29 | -0.20 | -0.51, 0.11 | 0.2 | 16 | -0.32 | -0.71, 0.08 | 0.12 |
| pure_bid_disperser | 70 | -0.36 | -0.59, -0.13 | 0.002 | 16 | -0.17 | -0.57, 0.22 | 0.4 |
| pure_gav_disperser | 42 | -0.10 | -0.37, 0.16 | 0.4 | 6 | -0.14 | -0.75, 0.46 | 0.6 |
| pure_niv_disperser | 24 | -0.42 | -0.78, -0.05 | 0.025 | 6 | -0.22 | -0.85, 0.42 | 0.5 |
| **Zero-inflation** | 463 | -8.019 | -8.042, 7.996 | >0.9 | 195 | -10.72 | -10,75, 10,70 | >0.9 |
| *^1^*CI = Confidence Interval | | | | | | | | |

**Table S9. Summary of generalized linear models (GLMs) testing effects of origin and origin categories on reproductive success in successful breeders male (n = 149) and female (n = 210) Atlantic salmon adults.** Poisson models tested effects of origin (0 = local; 1 = immigrant) and origin categories (Local_pur = Local, Local_without_parents = Local_Unrelated, Descendant_immigrant = Immigrant-F1, Hatchery_disperser = Immigrant-HAT, Immigrant-BID, Immigrant-GAV, Immigrant-NIV) on reproductive success. An offset term based on the logarithm of offspring sampled in each parent’s sampling year was included in the model to control for annual variation in sampling effort. N = sample size, IRR = Incidence Rate Ratio, 95% CI = 95% Confidence Interval, P = p-value. Reference category: 0 (Local) ; Local_pur (Local).

Table S9.1 – Origin and RS

| **Term** | **Parameter estimate** | **95% CI***^1^* | **P** |
| --- | --- | --- | --- |
| (Intercept) | -3.3 | -3.3, -3.2 | <0.001 |
| Origin |  |  |  |
| 0 | — | — |  |
| 1 | -0.15 | -0.24, -0.07 | <0.001 |
| *^1^*CI = Confidence Interval | | | |

Table S9.2 – Separated by sex: origin categories and RS

|  | **RS / FEMALES / BREEDERS** | | | | **RS / MALES / BREEDERS** | | | |
| --- | --- | --- | --- | --- | --- | --- | --- | --- |
| **Term** | **N** | **Parameter estimate** | **95% CI***^1^* | **P** | **N** | **Parameter estimate** | **95% CI***^1^* | **P** |
| **Intercept** | 210 | -3.1 | -3.2, -3.0 | <0.001 | 149 | -3.4 | -3.5, -3.3 | <0.001 |
| **Categorie** |  |  |  |  |  |  |  |  |
| local_pur | 87 | — | — |  | 48 | — | — |  |
| local_without_parents | 62 | -0.20 | -0.31, -0.08 | <0.001 | 40 | -0.07 | -0.23, 0.09 | 0.4 |
| descendant_immigrant | 14 | -0.59 | -0.81, -0.36 | <0.001 | 8 | 0.10 | -0.17, 0.37 | 0.5 |
| hatchery_disperser | 11 | 0.14 | -0.05, 0.34 | 0.15 | 13 | -0.35 | -0.60, -0.09 | 0.007 |
| pure_bid_disperser | 22 | -0.10 | -0.26, 0.06 | 0.2 | 18 | -0.48 | -0.72, -0.24 | <0.001 |
| pure_gav_disperser | 9 | -0.02 | -0.25, 0.20 | 0.8 | 14 | 0.01 | -0.21, 0.22 | >0.9 |
| pure_niv_disperser | 5 | -0.38 | -0.71, -0.05 | 0.025 | 8 | -1.1 | -1.6, -0.67 | <0.001 |
| *^1^*CI = Confidence Interval | | | | | | | | |

**Table S10. Summary of zero‐inflated Poisson / Poisson generalized linear models testing effects of number of partners, sex, origin, and their interactions on reproductive and mating success in Atlantic salmon adults (n=658) and successful breeders (n = 359).** Table S10.1 – Testing effect of sex (Female, n = 361; Male, n = 297), mating success and their interaction on RS. Table S10.2 – Testing effect of sex, origin, and mating success and their interactions on RS. Table S10.3 – Only successful breeders: Testing effect of sex (Female, n = 210; Male, n = 149), origin, and mating success and their interactions on RS. An offset term based on the logarithm of offspring sampled in each parent’s sampling year was included in both model components to control for annual variation in sampling effort. N = sample size, IRR = Incidence Rate Ratio, 95% CI = 95% Confidence Interval, P = p-value. Reference categories: Female, Local (origin = 0), and baseline number of partners.

Table S10.1 – Sex/Mating success and RS

| **Term** | **N** | **Parameter estimate** | **95% CI***^1^* | **P** |
| --- | --- | --- | --- | --- |
| **(Intercept)** | 658 | -4.1 | -4.2, -4.0 | <0.001 |
| **Number of partners** | 658 | 0.31 | 0.29, 0.33 | <0.001 |
| **Sex** |  |  |  |  |
| F | 361 | — | — |  |
| M | 297 | -0.73 | -0.91, -0.55 | <0.001 |
| **Number of partners * Sex** | 658 |  |  |  |
| Number of partners * M | 297 | 0.14 | 0.09, 0.19 | <0.001 |
| **Zero-inflation** | 658 | -0.51 | -0.70, -0.31 | <0.001 |
| *^1^*CI = Confidence Interval | | | | |

Table S10.2 – Sex/origin and Mating success

| **Term** | **N** | **Parameter estimate** | **95% CI***^1^* | **P** |
| --- | --- | --- | --- | --- |
| **(Intercept)** | 658 | -4.1 | -4.2, -4.0 | <0.001 |
| **Number of partners** | 658 | 0.30 | 0.27, 0.32 | <0.001 |
| **Sex** |  |  |  |  |
| F | 361 | — | — |  |
| M | 297 | -0.42 | -0.64, -0.21 | <0.001 |
| **Origin** |  |  |  |  |
| 0 | 449 | — | — |  |
| 1 | 209 | -0.17 | -0.42, 0.07 | 0.2 |
| **Number of partners * Sex** | 658 |  |  |  |
| Number of partners * M | 297 | 0.08 | 0.03, 0.14 | 0.004 |
| **Number of partners * Origin** | 658 |  |  |  |
| Number of partners * 1 | 209 | 0.07 | 0.01, 0.13 | 0.019 |
| **Sex * Origin** | 658 |  |  |  |
| M * 1 | 110 | -0.80 | -1.2, -0.40 | <0.001 |
| **Number of partners * Sex * Origin** | 658 |  |  |  |
| Number of partners * M * 1 | 110 | 0.13 | 0.03, 0.24 | 0.009 |
| **Zero-inflation** | 658 | -0.52 | -0.71, -0.33 | <0.001 |
| *^1^*CI = Confidence Interval | | | | |

Table S10.3 – Only successful breeders: Sex, origin and Mating success

| **Term** | **N** | **Parameter estimate** | **95% CI***^1^* | **P** |
| --- | --- | --- | --- | --- |
| **(Intercept)** | 359 | -4.1 | -4.2, -4.0 | <0.001 |
| **Number of partners** | 359 | 0.28 | 0.26, 0.31 | <0.001 |
| **Sex** |  |  |  |  |
| F | 210 | — | — |  |
| M | 149 | -0.26 | -0.45, -0.06 | 0.010 |
| **Origin** |  |  |  |  |
| 0 | 259 | — | — |  |
| 1 | 100 | -0.09 | -0.31, 0.13 | 0.4 |
| **Number of partners * Sex** | 359 |  |  |  |
| Number of partners * M | 149 | 0.04 | -0.01, 0.09 | 0.2 |
| **Number of partners * Origin** | 359 |  |  |  |
| Number of partners * 1 | 100 | 0.05 | 0.00, 0.11 | 0.063 |
| **Sex * Origin** | 359 |  |  |  |
| M * 1 | 53 | -0.73 | -1.1, -0.35 | <0.001 |
| **Number of partners * Sex * Origin** | 359 |  |  |  |
| Number of partners * M * 1 | 53 | 0.12 | 0.02, 0.22 | 0.019 |
| *^1^*CI = Confidence Interval | | | | |
